# Mechanisms underlying the defective development of xyloglucan-deficient root hairs in *Arabidopsis thaliana*

**DOI:** 10.64898/2026.10.13.758886

**Authors:** Vyankatesh Zambare, Gavin McDermond, Luis Alonso Baez, Hojae Yi, Thorsten Hamann, Charles T. Anderson

**Author notes:** Corresponding author: Charles T. Anderson, Department of Biology, The Pennsylvania State University, University Park, PA 16802 USA. Children’s Hospital of Philadelphia, Philadelphia, PA, 19104 USA. The author responsible for distribution of materials integral to the findings presented in this article in accordance with the policy described in the Author guidelines (https://academic.oup.com/plcell/pages/author-guidelines) is Charles T. Anderson.

## Abstract

Hemicellulosic xyloglucan is abundant in the primary cell walls of eudicots and non-gramineous monocots and contributes to their mechanical integrity. Its deficiency results in morphologically abnormal root hairs in Arabidopsis mutants lacking either two *XYLOGLUCAN XYLOSYLTRANSFERASE* (*xxt1 xxt2*) or five *CELLULOSE SYNTHASE-LIKE C* (*cslc-Δ5*) genes, but the cellular and biomechanical mechanisms underlying these abnormalities have remained unexplored. We combined multiple imaging modalities and quantitative modeling to unmask these mechanisms, first observing that xyloglucan-deficient mutants produced fewer and shorter root hairs that grew slower than wild type. Xyloglucan-deficient trichoblasts showed root hair initiation defects, helping to explain the sparseness of root hairs in these mutants. The mutant root hairs also had lower turgor pressure, thicker walls, and wider cross-sections, that explain lack of anisotropic growth, but cellulose organization appeared normal in xyloglucan-deficient root hairs. Brillouin microscopy suggested that xyloglucan-deficient root hairs were softer than wild-type root hairs. Finite element modeling of mutant root hairs predicted lower surface stress and growth rates, which were directly proportional to turgor pressure and inversely proportional to wall thickness. Stress patterns on models hint at potential mechanisms of bulging in xyloglucan-deficient root hairs. We propose that though cellulose orientation specifies the axis of cell expansion, it cannot enforce it without xyloglucan that interlinks microfibrils into a load-bearing network, identifying matrix-cellulose interaction as a distinct and necessary determinant of anisotropic plant cell growth.

## Introduction

In plants, root hairs enhance water and nutrient absorption by increasing the surface area of the primary root epidermis (Grierson et al., 2014). Root hairs also aid soil penetration by anchoring the root in heterogenous and compacted soils, as seen in maize and rice (Bengough et al., 2016; Kong et al., 2024). Root hairs have complex cell walls that modulate their biomechanical properties and influence their growth (Zambare & Anderson, 2026). Engineering crops with deeper roots is becoming important due to increasing cases of soil compaction and hardening, as deeper roots can potentially enhance water and nutrient absorption and hence crop productivity (Thorup-Kristensen et al., 2020).

Root hairs are encased in primary cell walls that are composed of cellulose, pectins, and hemicelluloses. Each type of polysaccharide contributes to the biomechanical properties of the wall depending on the growth stage of the root hair; for example, both pectin crosslinking and cellulose orientation can influence wall mechanics (Kerstens & Verbelen, 2003; Schoenaers et al., 2018). Hemicelluloses are a unique class of polysaccharides that contain a backbone with β-1,4 linkages in an equatorial configuration and can possess various side chains (Yoshimi et al., 2024). These polymers can interact with cellulose and influence wall mechanics (Cosgrove, 2016).

Xyloglucan is the most abundant hemicellulose in the primary walls of most eudicots and non-graminaceous monocots or type-I cell walls (Hayashi, 1989). It is composed of a β-1,4 glucan backbone and various sugar sidechains as decorations depending on the type of xyloglucan. The sidechains mainly consist of D-xylosyl groups that can link to D-galactosyl groups, which in turn can link to L-fucosyl groups (Faik et al., 2002). Xyloglucan is synthesized in the Golgi apparatus before it is transported to the cell surface and incorporated into the cell wall (Kim et al., 2020).

In *Arabidopsis thaliana*, *CELLULOSE SYNTHASE LIKE-C* (*CSLC*) genes are responsible for the synthesis of the xyloglucan backbone, whereas XYLOGLUCAN XYLOSYLTRANSFERASES (XXTs) attach xylose sidechains to the backbone (Cavalier & Keegstra, 2006; Kim et al., 2020). Previous studies have shown that the *cslc-Δ5* quintuple mutant that lacks all five paralogous *CSLC* genes (*CLSC4, CSLC5, CSLC6, CSLC8, and CSLC12)*, and the *xxt1 xxt2* double mutant, lack any detectable xyloglucan and therefore serve as excellent tools to study the effects of xyloglucan deficiency on the cellular and physiological properties of plants (Bou Daher et al., 2024; Cavalier et al., 2008; Kim et al., 2020).

Xyloglucan deficiency also affects cellular and tissue mechanics and mechanosensitivity (Codjoe et al., 2022; Jonsson et al., 2022; Xiao et al., 2016). Etiolated hypocotyls of *xxt1 xxt2* display an altered three-dimensional distribution of cellulose, which is the major load-bearing polymer in plant cell walls, and also show reduced stability of microtubules against external mechanical pressure (Xiao et al., 2016). In the case of *cslc-Δ5*, etiolated hypocotyls show reduction in cellulose crystallinity and turgor pressure, in addition to softer and weaker cell walls (Bou Daher et al., 2024). Petioles of *xxt1 xxt2* show more mechanical compliance, but undergo less expansin-mediate creep than wild-type petioles (Park & Cosgrove, 2012), which suggests that xyloglucan is involved in reinforcing cell walls as well as in α-expansin-mediated wall loosening. The *cslc-Δ5* mutant also shows fewer pollen tubes after pollination, which suggests a function for xyloglucan in polar growth in plant cells (Kim et al., 2020). Previous studies have shown that xyloglucan-deficient mutants have more bulged root hairs with uneven shapes, but the cellular and molecular mechanisms underlying this defective growth are still unclear (Cavalier et al., 2008; Zabotina et al., 2012). Therefore, in this study, we sought to mechanistically link the cellular phenotypes and biomechanical properties of xyloglucan-deficient root hairs in *Arabidopsis thaliana* mutants *xxt1 xxt2* and *cslc-Δ5* to the observed growth defects.

We quantified the phenotypes of xyloglucan-deficient root hairs and observed that they produce fewer and shorter root hairs that grow slower than wild-type root hairs. The trichoblasts of mutants were longer than those of the wild type and showed fewer root hair bulges than those in wild type, indicating a possibility of defective root hair initiation. In both mutants under study, root hairs also had larger cross-sectional areas than wild type and thicker walls with lower turgor pressure. We further estimated mechanical properties such as elastic modulus (stiffness) and loss modulus (viscosity) in these root hairs using Brillouin microscopy and observed that xyloglucan-deficient root hairs showed less stiff behavior but no changes in their viscous behavior compared to wild type depending on growth stage and region measured. Lastly, we simulated the growth of root hairs using Finite Element Modeling (FEM) to study wall stress patterns in wild type and xyloglucan-deficient root hairs to infer the effects of cellular and mechanical properties on their phenotypes. Taken together, our data provide quantitative evidence to explain the defective development of root hairs in xyloglucan-deficient mutants of Arabidopsis and can be used to predict the growth patterns of root hairs in other genotypes and plant species.

## Results

### Xyloglucan deficiency results in shorter and fewer root hairs that grow slower

Xyloglucan-deficient root hairs have been reported to show varying morphological abnormalities in *Arabidopsis thaliana* (Cavalier et al., 2008; Kim et al., 2020; Zabotina et al., 2012). To explore these phenotypes in detail we quantified root hair length and density in *xxt1 xxt2* and *cslc-Δ5* mutants, which lack the side chains and backbone of xyloglucan, respectively. Qualitative observations of 5-day-old seedlings indicated that xyloglucan deficiency results in shorter and fewer root hairs and also shorter primary roots in *xxt1 xxt2* **(Fig. 1A)**. Upon quantification, we found that *xxt1 xxt2* roots with an average length of 10.89 ± 2.56 (mean ± SD) mm were shorter than Col-0 roots with an average length of 14.07 ± 2.63 mm **(Fig. 1B)**, whereas *cslc-Δ5* roots with an average length of 12.84 ± 2.18 mm were not significantly shorter than Col-0. Although there was a difference in root lengths between the mutants, roots were consistently wider in the maturation zones of both *xxt1 xxt2* (124.8 ± 9.07 µm) and *cslc-Δ5* (129.1 ± 5.66 µm) seedlings than in Col-0 (117.5 ± 7.85 µm) seedlings **(Fig. 1C)**, consistent with the wider diameter of etiolated hypocotyls in the *xxt1 xxt2* mutant (Xiao et al., 2016). Wider roots could potentially provide more surface area for additional files of epidermal cells and thus form more trichoblasts as long as cell diameter does not vary significantly. This led us to hypothesize that more root hairs would be present in both mutants. However, both *xxt1 xxt2* and *cslc-Δ5* seedlings had significantly fewer root hairs per root (122.8 ± 56.68 and 128.3 ± 59.99, respectively) than Col-0 seedlings (236.0 ± 84.20 root hairs per root) **(Fig 1D)**. Fewer root hairs can arise due to shorter roots in mutants **(Fig. 1A),** hence root hair density provides more accurate quantification of root hair population. We found lower root hair density (root hairs per mm of root length) in both *xxt1 xxt2* (11.37 ± 5.08 root hairs/mm) and *cslc-Δ5* (10.22 ± 4.94 root hairs/mm) compared to Col-0 (16.59 ± 4.69 root hairs/mm) **(Fig 1E)**.

**Fig 1.**
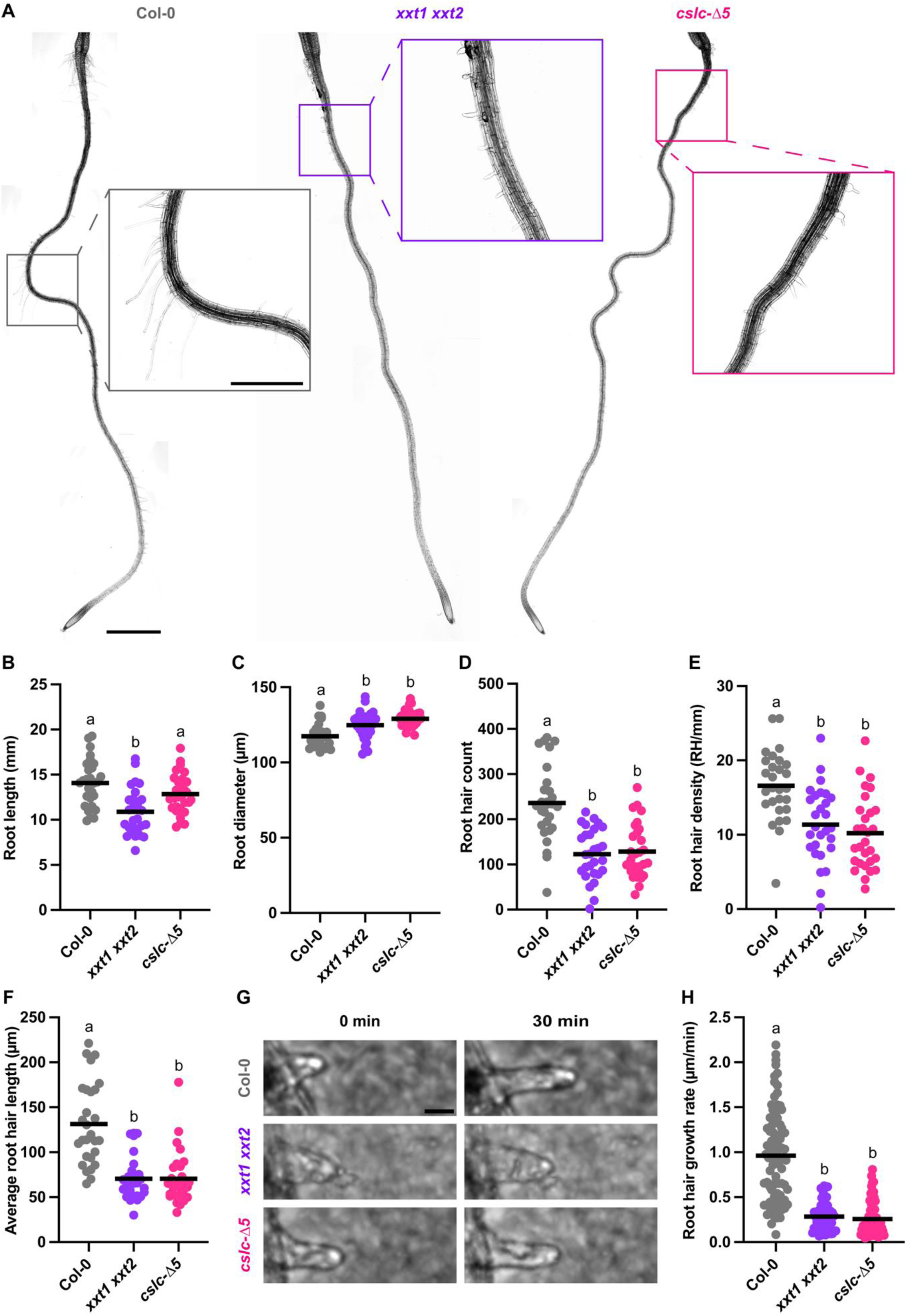
Root hair density, length, and growth rate are diminished in xyloglucan-deficient mutants. **A,** Representative images of 5-day old seedlings of Col-0, *xxt1 xxt2*, and *cslc-Δ5* grown on media plates (Scale bar = 1 mm, applies to all the panels). Insets show digitally zoomed-in roots for better observation of root hairs (Scale bar = 500 µm, applies to all the insets). **B,** Root length. **C,** Root diameter measured at least at three locations in the late maturation zone of root and averaged per root. **D,** Root hair count per seedling. **E,** Root hair density represented by the number of root hairs per mm of root length. Each point is one seedling (n = 28-29 seedlings per genotype). **F,** Average root hair length, measured on five root hairs per seedling in the late maturation zone. **G,** Representative brightfield images of root hairs growing in macrofluidic chambers (Scale bar = 20 µm). **J,** Root hair growth was imaged for 30 minutes to quantify growth rate. Each point represents one root hair (n = 50-85 root hairs from at least 15 seedlings per genotype). Horizontal lines represent averages Letters represent statistically separable groups, p < 0.05, one-way ANOVA and post-hoc Tukey test.

On average, root hairs in both xyloglucan mutants, *xxt1 xxt2* (70.44 ± 24.89 µm) and *cslc-Δ5* (70.59 ± 29.33 µm), were significantly shorter than Col-0 (131.5 ± 45.76 µm) **(Fig. 1F)**. To determine whether the shorter root hairs we observed in *xxt1 xxt2* and *cslc-*Δ*5* arose from differences in growth rate or duration, we performed timelapse imaging of actively growing root hairs into granular agar-based media in roots cultured in macrofluidic chambers. Unlike wild-type root hairs growing into stiffer media that grow faster but arrest their growth sooner than in softer media (Zambare et al., 2026), xyloglucan-deficient root hairs grew more slowly than wild-type: growth rates in *xxt1 xxt2* and *cslc-Δ5* were 0.28 ± 0.16 µm/min and 0.26 ± 0.19 µm/min, respectively, whereas Col-0 root hairs grew at a rate of 0.96 ± 0.50 µm/min **(Fig. 1G and H)**.

To explore whether the shorter root hairs we observed in *xxt1 xxt2* and *cslc-Δ5* arise from premature growth arrest, we divided the average final root hair lengths by the observed growth rate, estimating growth durations of 252 min, 272 min, and 136 min for *xxt1 xxt2*, *cslc-Δ5*, and Col-0, respectively. This estimate implies that xyloglucan-deficient root hairs do not arrest their growth prematurely, although it is possible that the growth rates we measured are not uniform across the entire growth period. Together, these data indicate that a lack of xyloglucan inhibits root hair development by leading to slower growth and potential root hair initiation defects.

### Xyloglucan-deficient trichoblasts are longer than wild-type trichoblasts and occasionally fail to initiate root hairs

Trichoblast length and root hair development can be linked through density and growth tradeoffs. In a study of redox control of root hair morphogenesis in Arabidopsis, it was found that trichoblast length and root hair density are inversely proportional (Sánchez-Fernández et al., 1997), and wild-type trichoblasts are shorter than atrichoblasts, even before the expected appearance of root hairs (Beemster & Baskin, 1998). In addition, in barley (*Hordeum vulgare*) asymmetric cell divisions precede the formation of root hairs, with the shorter cell becoming the trichoblast, and the *root hairless 1.b* (*rhl1.b*) mutant that lacks root hairs only has a single type of longer epidermal cell (Marzec et al., 2013). It is possible that in Arabidopsis, trichoblasts, with regards to dedicating cellular machinery and metabolites to root hair production, do not sustain longitudinal growth as highly as atrichoblasts. Therefore, based on the shorter root hairs and decreased root hair density phenotypes **(Figs. 1E & F)**, we hypothesized that both *xxt1 xxt2* and *cslc-Δ5* mutants have longer trichoblasts.

As expected, we found that cells in trichoblast files in *xxt1 xxt2* (165.60 ± 22.44 µm) and *cslc-Δ5*(179.90 ± 25.06 µm) were longer than the equivalent cells in Col-0 (150.70 ± 24.12 µm) **(Fig. 2A & B)**. Cells in atrichoblast files in *xxt1 xxt2* (208.28 ± 29.45 µm) and *cslc-Δ5* (220.86 ± 26.76 µm) were similar in length to the equivalent cells in Col-0 (211.88 ± 22.10 µm) **(Supp. Fig. 2C)**.

**Fig 2.**
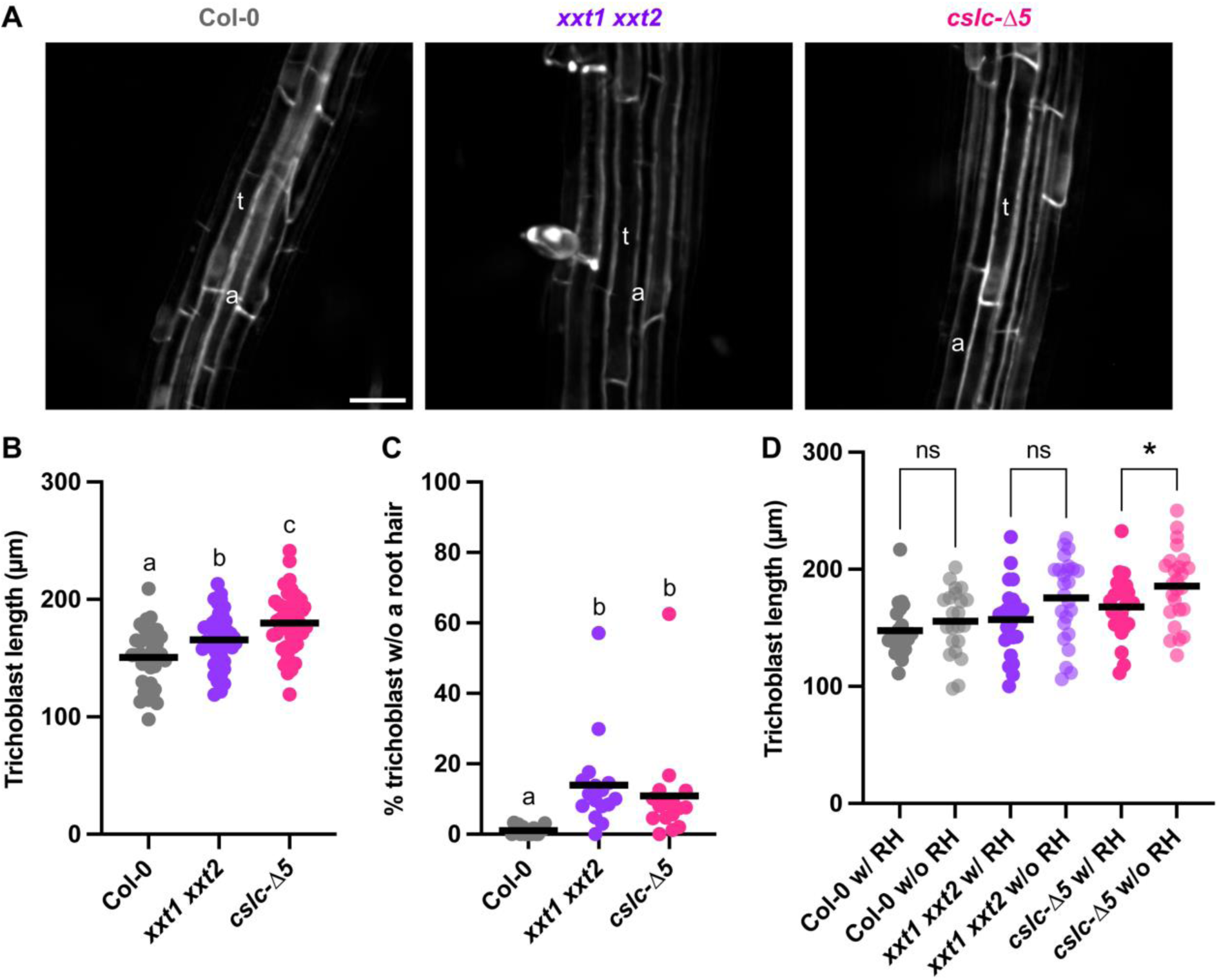
Xyloglucan deficient mutants have longer trichoblasts and show higher rates of defective root hair initiation. **A,** Images represent maturation zones of 5-day old seedlings of Col-0, *xxt1 xxt2*, and *cslc-Δ5*, t = trichoblast, a = atrichoblast (Scale Bar = 50 µm, applies to all the panels). **B,** Trichoblast lengths averaged from at least three epidermal cells in the maturation zone of roots (n = 38-46 seedlings per genotype). **C,** Percentage of trichoblast population (n ≥ 50 cells per root) in the maturation zone without any root hair or even a root hair bulge (n = 16 seedlings per genotype). **D,** Lengths of trichoblasts with and without a root hair (RH; n = 22-25 seedlings per genotype, 1-11 trichoblasts/seedling). Horizontal lines represent averages. Each data point represents one seedling. In B and C, different letters above the graph represent statistically significant difference compared to the control, p < 0.05, one-way ANOVA and post-hoc Tukey test. For D, Student’s t-test, p < 0.05, was used to compare the means of trichoblasts with and without root hairs within individual genotypes; genotypes were not compared to one another.

Finding the percentage of trichoblasts that do not form root hairs would give a clear idea whether the decreased root hair density we observed in mutants is solely because of the longer trichoblasts, or because a proportion of trichoblasts fail to initiate root hairs. We observed that the percentage of trichoblast that did not form root hairs in *xxt1 xxt2* (14 ± 13.36 %) and *cslc-Δ5* (10.87 ± 14.46 %) was higher when compared to Col-0 (1.06 ± 1.25 %) **(Fig. 2C)**. We also estimated the number of trichoblasts in a single cell file along the length of the root using average values of root length **(Fig. 1B)** and root hair count **(Fig. 1D)** and found that Col-0 has 93.36 trichoblasts on average which was more than *xxt1 xxt2* and *cslc-Δ5* which had 65.76 and 71.37 trichoblasts per root respectively. These trichoblast counts per root are underestimated and mainly reflect the number of trichoblasts in a single cell file but still give a decent estimate for the amount of trichoblasts present in a single root. These results suggest that the decreased root hair density observed in xyloglucan-deficient mutants arises because of the longer and hence potentially sparser trichoblasts in their roots, that sometimes fail to initiate root hairs.

We also studied epidermal cell patterning in roots and though we observed wider roots in xyloglucan-deficient mutants **(Fig. 1C)**, there were no changes in the proportions of trichoblasts and atrichoblasts along the root girth across genotypes **(Supp. Fig. 1)**. This suggests that root hair density is not explained by the proportion of trichoblasts along the root girth as was the case when Col-0 roots experienced increasing mechanical impedance and show fewer proportion of trichoblasts and hence fewer root hairs (Zambare et al., 2026).

To test the resource allocation hypothesis stated earlier, i.e, whether there is a relation between the longitudinal growth of trichoblasts and root hair morphogenesis, we quantified the lengths of trichoblasts with and without root hairs and compared within genotypes **(Fig. 2C)**. There was no significant difference between the trichoblasts with and without a root hair in Col-0 and *xxt1 xxt2* roots, but the trichoblasts without a root hair in *cslc-Δ5* roots were significantly longer than those with a root hair. We also compared the lengths of trichoblasts with and without a root hair between different genotypes and found that the trichoblasts with a root hair in *cslc-Δ5* roots (167.90 ± 26.11 µm) were significantly longer than those in Col-0 (147.60 ± 22.49 µm) and *xxt1 xxt2* (157.20 ± 29.75 µm) **(Supp. Fig. 2A)**. A similar trend was observed in the trichoblasts without a root hair as in *cslc-Δ5* (185.60 ± 31.70 µm) they were significantly longer than in Col-0 (155.60 ± 28.08 µm) and *xxt1 xxt2* (175.70 ± 35.19 µm) **(Supp. Fig. 2B)**.

In *xxt1 xxt2* and *cslc*-Δ5 roots, the differences between mean lengths of the trichoblasts with and without a root hair were 18.5 µm and 17.7 µm, respectively. Though the means in *xxt1 xxt2* are not statistically separable, the biological trend is quite comparable to *cslc-Δ5*. Therefore, in xyloglucan-deficient mutants, the data support a model in which cells in trichoblast files make tradeoffs between longitudinal growth and root hair production. However, the chronological control between root hair morphogenesis and trichoblast elongation is still unclear.

### Root hairs are wider, and their cell walls are thicker, in xyloglucan-deficient mutants

We hypothesized that anisotropic growth in root hairs is influenced by cell wall thickness and the orientation of cellulose fibrils. Thicker cell walls in the shank might favor tip growth. By contrast cellulose having a high tensile modulus (defined as the resistance to elastic deformation) should favor wall expansion perpendicular to the main direction of fibril deposition (Baskin et al., 2004). When imaging root hairs in 3D, we observed that root hair cross-sections in xyloglucan-deficient mutants *xxt1 xxt2* (198.80 ± 81.86 µm^2^) and *cslc-Δ5* (202.15 ± 93.31 µm^2^) were wider than in Col-0 (60.22 ± 21.29 µm^2^) (Fig. 3A and B). The significantly shorter **(Fig. 1F)** and broader **(Fig. 3B)** root hairs in xyloglucan-deficient mutants indicate that their anisotropic expansion is perturbed. Therefore, we quantified cell wall thickness and found that walls in the shank of root hairs in *xxt1 xxt2* (548.09 ± 75.19 nm) and *cslc-Δ5* (572.31 ± 75.96 nm) seedlings were thicker than in Col-0 seedlings (465.75 ± 52.75 nm) **(Fig. 3C)**.

**Fig 3.**
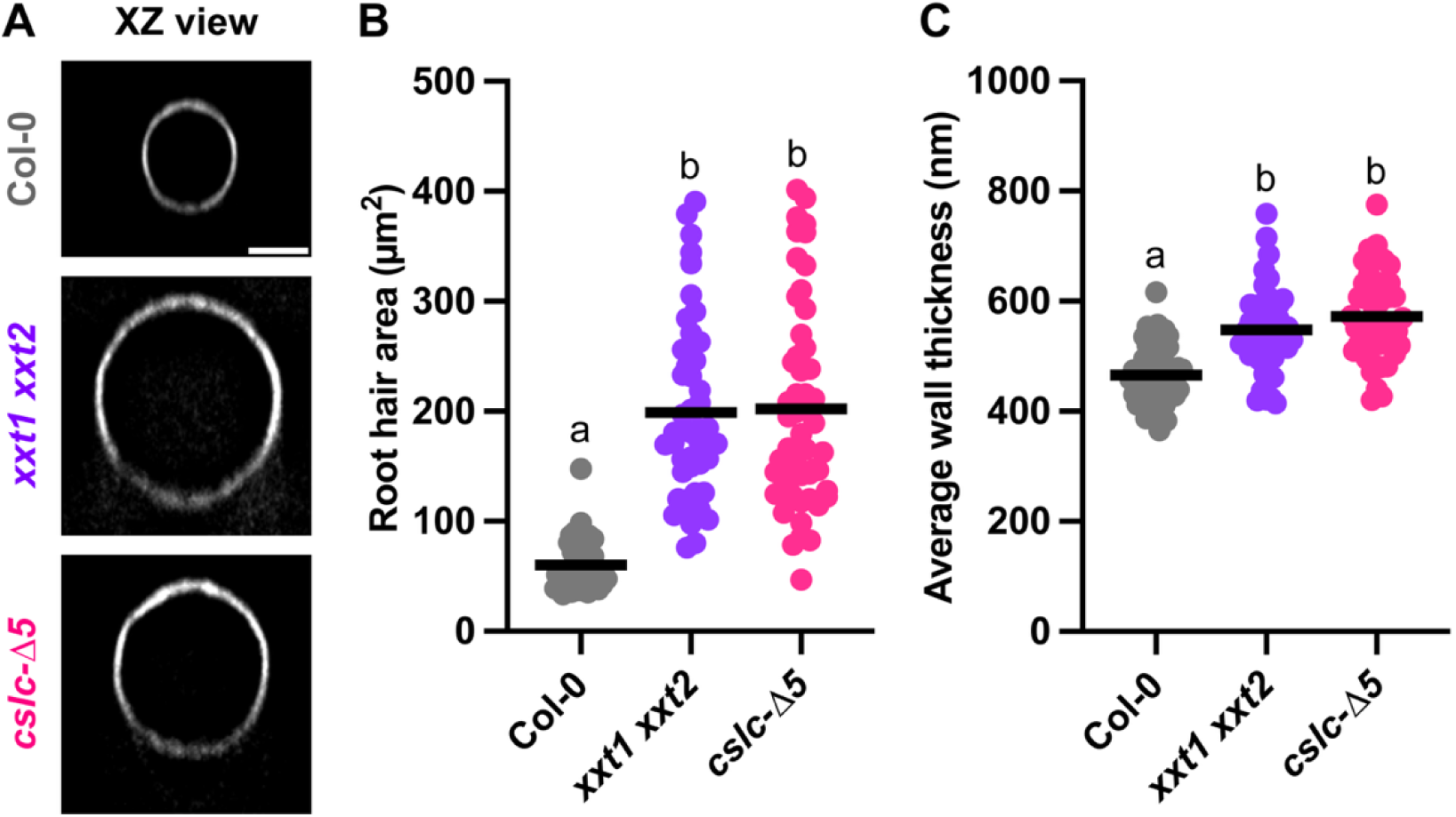
Xyloglucan deficiency increases root hair cross-sectional area and cell wall thickness. **A,** Images represent 5-day old seedlings of Col-0, *xxt1 xxt2*, and *cslc-Δ5* (Scale bar = 5 µm, applies to all the panels). Fluorescence signal represents S4B staining. **B,** XZ-cross-sectional area of root hairs averaged from at least three locations in root hair shanks. **C,** Average cell wall thickness along the root hair shank. Each point on the graph represents one root hair (n = 45-54 root hairs per genotype from at least 15 seedlings). Horizontal lines represent averages. Different letters above the graph represent statistically significant difference, p < 0.05, one-way ANOVA and post-hoc Tukey test.

Root hair diameter and perimeter also followed the same trend as the cross-sectional area of root hairs. In *xxt1 xxt2* and *cslc-Δ5*, the root hair diameters were 15.30 ± 3.12 µm and 15.55 ± 3.75 µm respectively, whereas in Col-0 the root hair diameter was 8.72 ± 1.28 µm **(Supp. Fig. 3A)**. The perimeters of the root hairs in *xxt1 xxt2* (48.85 ± 9.86 µm) and *cslc-Δ5* (49.24 ± 11.37 µm) were larger than in Col-0 (27.34 ± 4.53 µm) **(Supp. Fig. 3B)**. The cellulose orientation and optical anisotropy of cellulose fibrils in the root hairs of both the mutants, as quantified with the FibrilTool plug-in, were not significantly different than wild type **(Supp. Fig. 4)**. Therefore, the cellulose fibril orientation does not explain the isotropic growth in *xxt1 xxt2* and *cslc-Δ5* root hairs.

As stated earlier, thicker walls would be expected to promote anisotropic growth in root hairs, which is clearly not the case in both xyloglucan-deficient mutants. Therefore, we decided to quantify subcellular and wall mechanical properties to investigate their role in the abnormal root hair growth we observed.

### Xyloglucan-deficient trichoblasts have lower turgor pressure

An intricate balance is required between cell wall deposition, wall remodeling, and turgor pressure to facilitate plant cell expansion (Ali et al., 2023; Shackel et al., 1987). A turgor pressure above the yield threshold of cell wall is required for root hair elongation (Alline et al., 2026). In etiolated hypocotyls of Arabidopsis, xyloglucan deficiency led to a lower amount of crystalline cellulose and softer and weaker cell walls (Bou Daher et al., 2024). In the same study, the authors also observed lower turgor pressure in epidermal cells of etiolated hypocotyls by using plasmolysis and microextensometer-based mechanical measurements. Based on the literature and the root hair phenotypes we observed, we hypothesized a lower turgor pressure in xyloglucan-deficient trichoblasts. We used lines with fluorescently tagged markers of the plasma membrane and an incipient plasmolysis protocol described previously to quantify turgor pressure in wild type and xyloglucan-deficient roots (Zambare et al., 2026). Upon quantification, we observed that more trichoblasts plasmolyzed at the same sorbitol concentration in xyloglucan-deficient mutants **(Fig. 4A)**, which indicates that they have lower turgor pressure compared to their corresponding wild-type lines **(Fig. 4B).** Turgor pressures values were calculated as follows: LTI6b-GFP Col-0 (0.801 ± 0.028 MPa), LTI6b-GFP *xxt1 xxt2* (0.637 ± 0.054 MPa), MYR-YFP Col-0 (0.794 ± 0.053 MPa), and MYR-YFP *cslc-Δ5* (0.605 ± 0.138 MPa). This trend is similar to the one observed in etiolated hypocotyls of Arabidopsis (Bou Daher et al., 2024), and further explains the potential mechanisms behind shorter and fewer root hairs in xyloglucan-deficient mutants.

**Fig 4.**
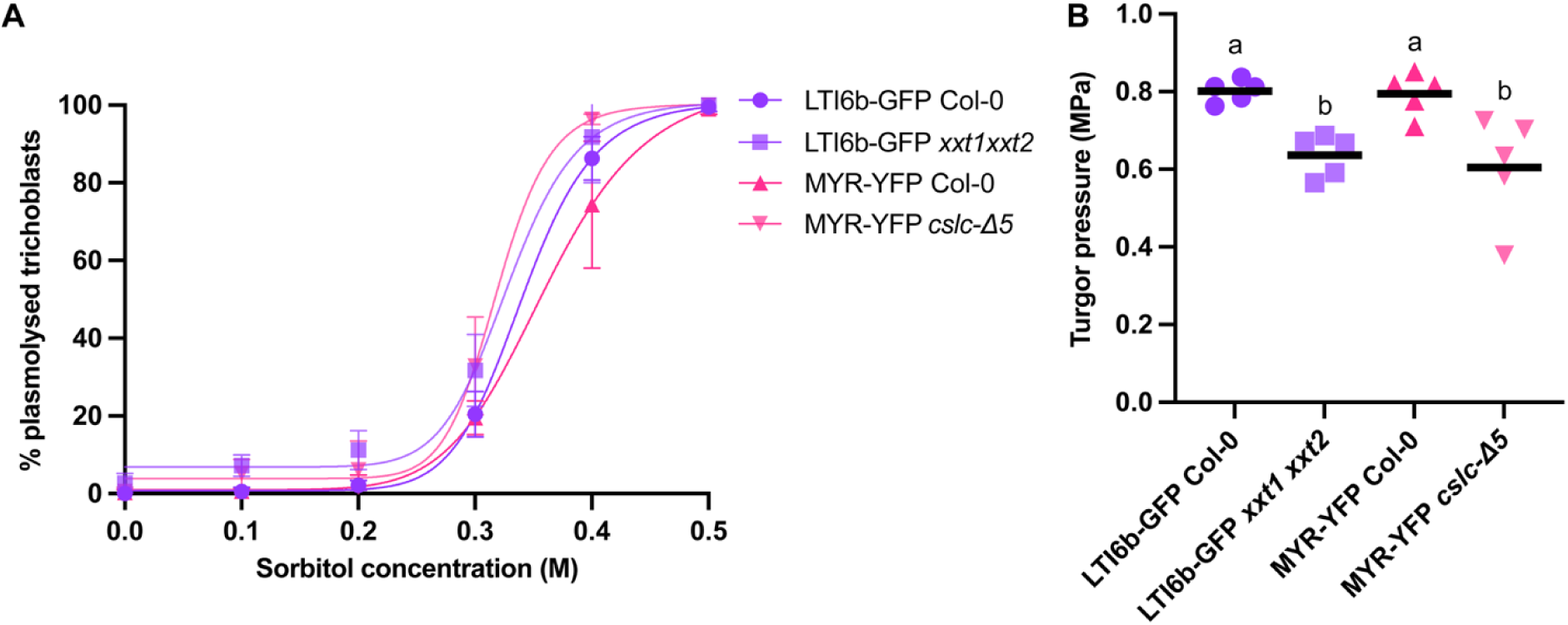
Xyloglucan-deficient trichoblasts have lower turgor pressure. **A,** Non**-**linear regression curves representing percentage of plasmolyzed trichoblasts in increasing concentrations of sorbitol. Error bars represent standard deviation. **B,** Turgor pressure as calculated using the plasmolysis trends. Each data point represents one independent experiment (n = 12-16 seedlings per genotype per sorbitol concentration). Horizontal lines represent averages. Different letters above the graph represent statistically significant difference compared to the control, p < 0.05, Student’s t-test was used to compare pairs of genotypes expressing the same plasma membrane marker.

### Xyloglucan-deficient root hairs are softer than wild type, depending on the growth stage and genotype

As we observed thicker walls but lower turgor pressures in xyloglucan-deficient root hairs, we hypothesized a decrease in the stiffness of these root hairs as it is modulated by turgor pressure in live cells (Beauzamy et al., 2014). We used Brillouin microscopy, a non-invasive optical technique, to investigate the mechanical properties of root hairs (Alonso Baez et al., 2026; Samalova et al., 2024). We imaged membrane tagged fluorescent maker lines of Col-0, *xxt1 xxt2,* and *cslc-Δ5* **(Fig. 5A & B, upper rows)** using the marker to locate the central plane of the root hair for imaging. The frequency shift of the Brillouin spectrum is correlated with the stiffness of the biological sample, whereas the linewidth gives an estimation of the viscosity of the sample (Prevedel et al., 2019). Using frequency shift data, we also estimated the storage modulus which measures the material’s ability to store elastic energy when it undergoes compressions or rarefactions at the molecular level in one direction without being allowed to expand in the other directions. The storage modulus is thus a measure of the material stiffness. The linewidth data was used to estimate the viscosity (loss modulus) of root hairs, which gives a measure of the mechanical energy lost through viscous relaxation during the deformation. These moduli describe the high-frequency (GHz range) mechanical behavior of a material by probing thermally generated acoustic waves through light scattering. This contrasts with the Young’s modulus, which describes the material’s low-frequency mechanical responses by applying a force and measuring deformation. We measured these parameters in root hair bulges, mature root hairs and on the borders/edges as well as on the area of root hairs, to differentiate mechanical contributions by the cell wall and the rest of the cell at distinct growth stages **(Table 1)**. The representative heatmaps of frequency shifts suggest that root hair bulges and mature root hairs might be softer in both xyloglucan-deficient mutants compared to their corresponding wild types as mutants had lower frequency shifts than wild-type root hair in both the border and area of the bulge and mature root hairs **(Fig. 5A & B, lower rows)**. However, only border values in the mature root hairs of *xxt1 xxt2* were significantly lower than wild type and in *cslc-Δ5* both border and area values in root hair bulges were lower than in their corresponding wild type **(Table 1)**.

**Fig 5.**
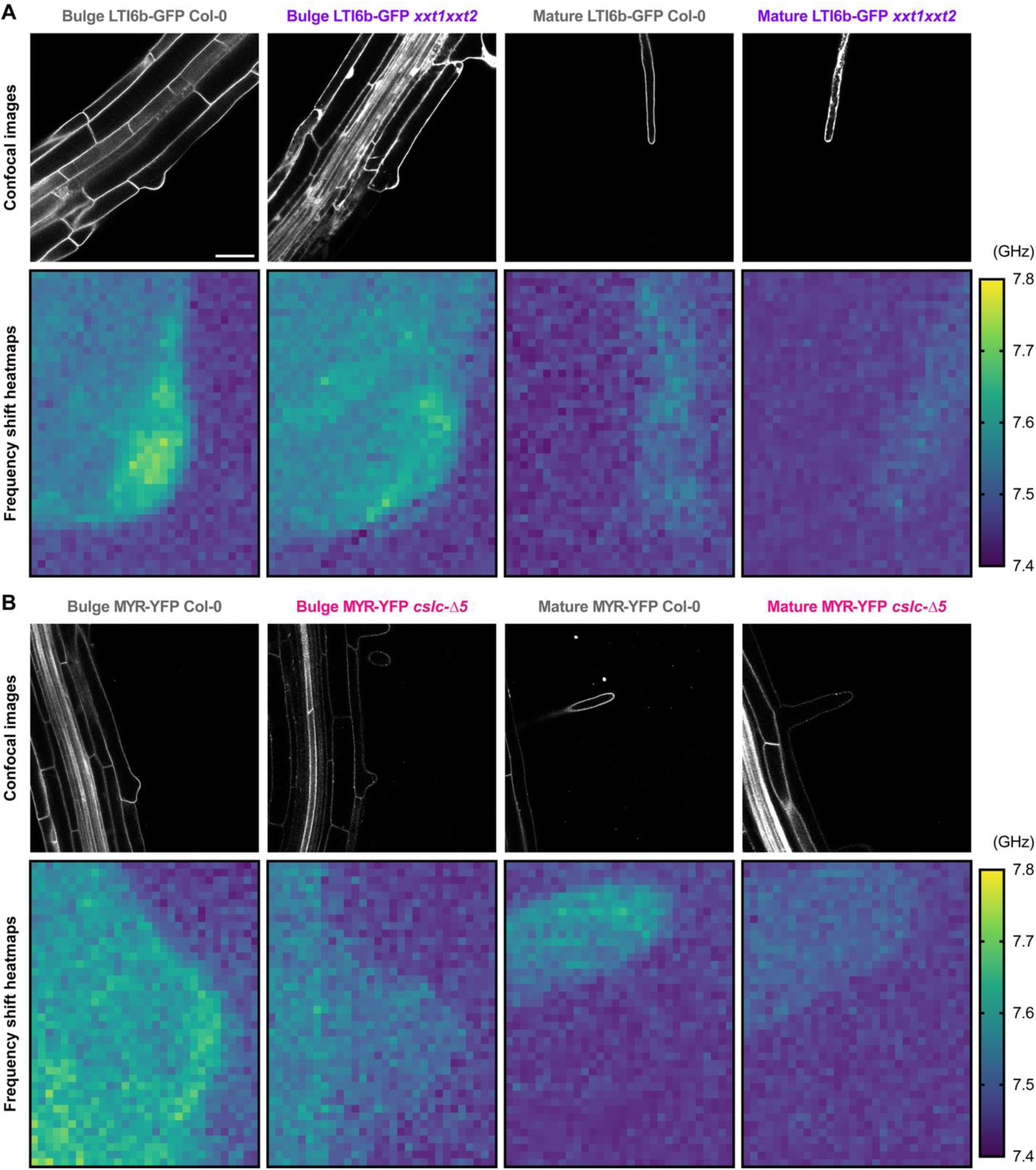
Brillouin microscopy reveals developmentally dependent lower stiffness in xyloglucan-deficient root hairs. **A, Upper row:** Confocal micrographs of root hair bulges and mature root hairs of Col-0 and *xxt1 xxt2* with their corresponding fluorescent tags highlighting the plasma membrane (Scale bar = 100 µm, applies to all the confocal panels), **Lower row:** Representative heatmaps showing frequency shift values of the Brillouin spectra acquired from corresponding root hairs. **B, Upper row:** Confocal micrographs of root hair bulges and mature root hairs of Col-0 and *cslc-Δ5* with their corresponding fluorescent tags highlighting the plasma membrane, **Lower row:** Representative heatmaps showing frequency shift values of the Brillouin spectra acquired from corresponding root hairs. n ≥ 10 root hairs from at least 10 different

**Table 1:**
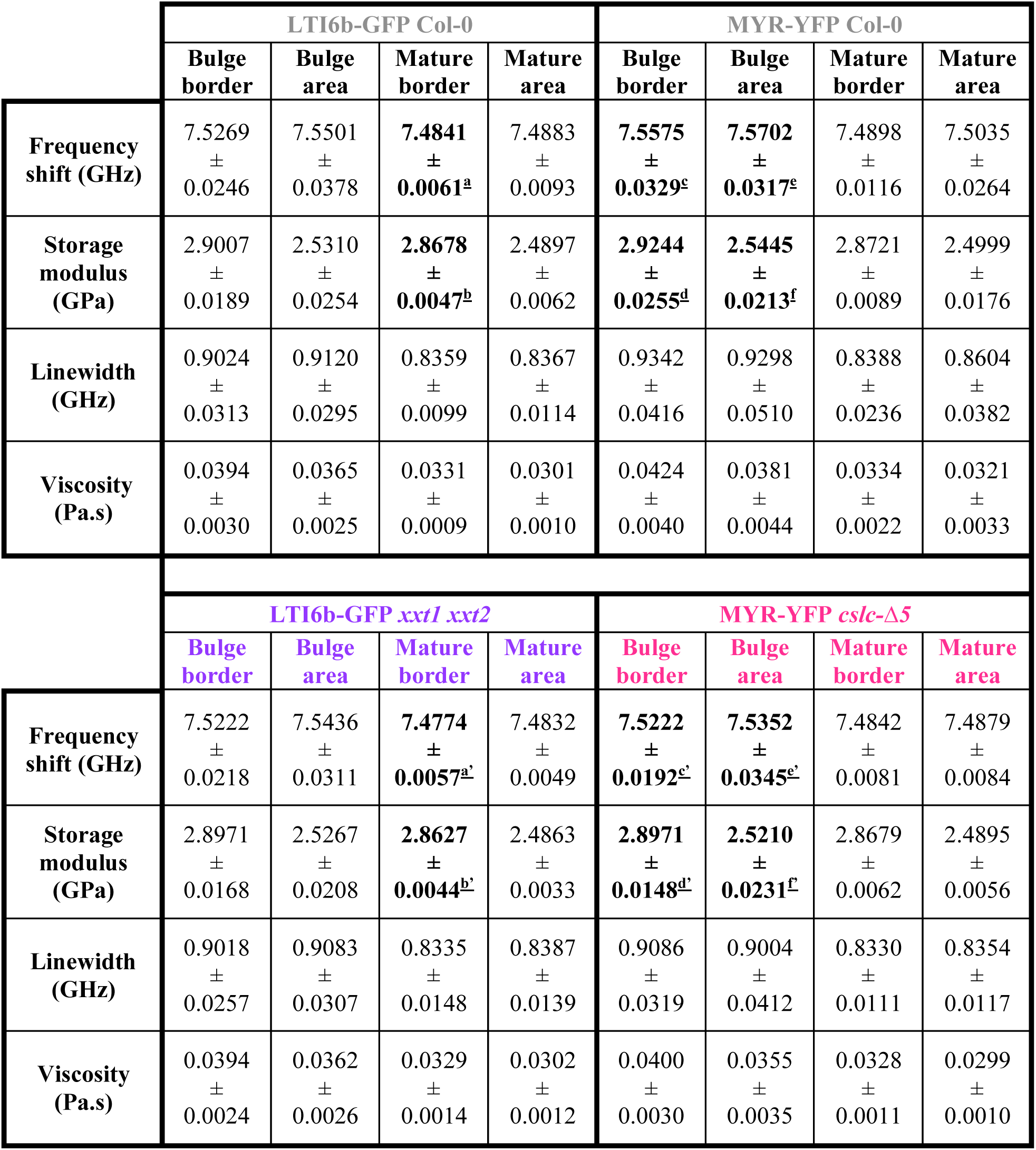
Estimation of mechanical properties of root hairs using Brillouin microscopy. Brillouin spectra give the magnitudes of the frequency shift and linewidth that are used to estimate storage modulus (stiffness) and viscosity (loss modulus) of the sample, respectively. Different letters in the superscript (bold) represent statistically significant differences compared to the controls, p < 0.05; student’s t-test was used to compare pairs of genotypes expressing the same plasma membrane marker. n ≥ 10 root hairs from at least ten different seedlings.

The linewidth and hence the loss modulus (viscosity) values of root hair bulges and mature root hairs of *xxt1 xxt2* and *cslc-Δ5* were lower than their corresponding wild type hairs, but the differences were not statistically separable **(Supp. Fig. 5) (Table 1)**. However, the trend toward decreased viscosity, which is measured perpendicular to the growth axis of the root hair, corresponds with the wider root hairs observed in both the xyloglucan-deficient mutants **(Fig. 3) (Supp. Fig. 5)**. Lower viscosity means higher fluidity of material along the transverse axis of root hairs, which might lead to isotropic growth and hence wider root hairs.

### Finite element models mimic the *in vivo* growth patterns of root hairs and reveal complex stress patterns in xyloglucan-deficient root hairs

Plant cell shape and its functions are interconnected, and mechanical stress often influences cell morphogenesis (Sapala et al., 2018). To study the stress patterns on the walls of root hairs, we simulated their growth computationally using FEM. We built separate models for Col-0, *xxt1 xxt2*, and *cslc-Δ5* that reflect their length, diameter, and aspect ratio as observed *in vivo*. For these models, we incorporated average wall thicknesses obtained from confocal microscopy **(Fig. 3C)** and individual values for turgor pressure observed within replicates for each genotype **(Fig. 4)**. An additional turgor pressure value was generated by averaging turgor pressures for each genotype. The modeling results **(Fig. 6)** revealed that the surface stress in the modeled root hairs was mainly influenced by the turgor pressure, therefore capturing this variation in the stress distribution as a function of changing turgor pressure would reveal nuances of root hair growth in wild-type and xyloglucan-deficient mutants **(Figs. 6A-C)**. Upon simulating root hair growth, we observed that Col-0 root hairs expanded faster than both mutants, as was observed experimentally **(Fig. 1)**. The simulated growth rate of root hairs in Col-0 was 1.53 ± 0.91 µm/min, whereas in *xxt1 xxt2* root hairs, it was 0.57 ± 0.04 µm/min, and in *cslc-Δ5* root hairs, it was 0.59 ± 0.12 µm/min **(Fig. 6D)**. In terms of surface stress, we observed a complex distribution of stress on the finite elements at the tip of the root hair, around the rest of the dome, and at the sub-apical region. Therefore, we extracted and plotted stress values from these individual regions **(Fig. 6E-G)**, and the actual magnitudes are available in **Supp. Table 1**. Irrespective of the regions, stress values on both the mutants were comparable and were significantly lower than wild type.

**Fig 6.**
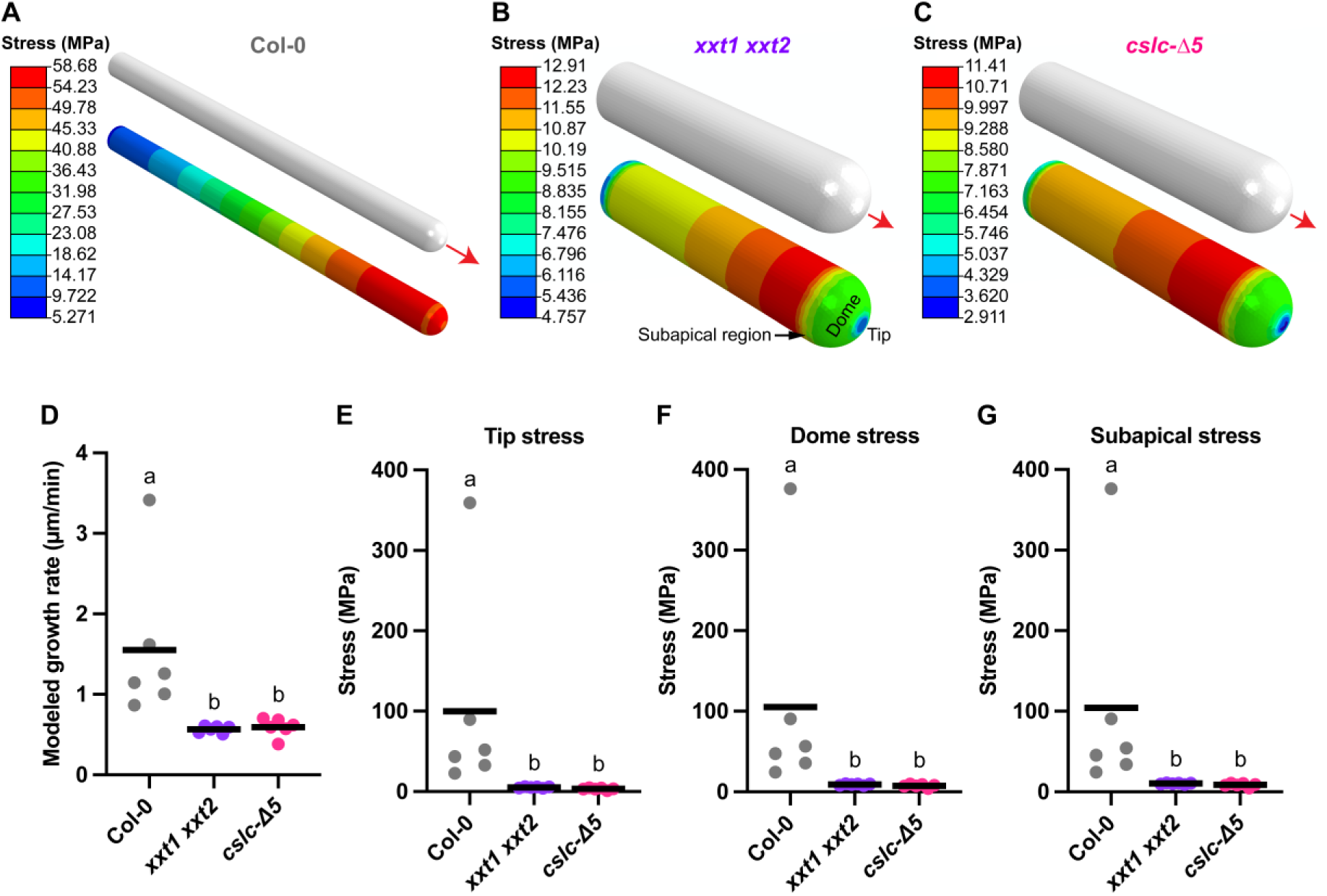
Finite element models accurately simulate root hair growth and help derive stress patterns on root hair surfaces. Models shown in this figure were run using the average values of turgor pressure for every genotype. **A-C,** Representative models of Col-0, *xxt1 xxt2*, and *cslc-Δ5*, before and after the run, respectively. Root hair tip, dome, and subapical regions are shown in **B** for representation. **Note:** Color scale range is different across panels A-C. **D,** Growth rate quantified by the displacement of finite elements before and after the run under unit time. **E-G,** Quantitation of stress patterns observed on the walls of root hairs *in silico* after each model run. Individual data points represent a single root hair or an experimental run. Horizontal lines represent averages. n = 6 root hairs per genotype. Letters represent statical significance, p < 0.05, one way ANOVA with post-hoc Tukey test.

The surface stress on FEMs is directly proportional to the turgor pressure and inversely proportional to the wall thickness as observed by swapping these parameters across genotypes **(Supp. Tables 2 and 3)**. The variability in stress patterning and higher surface stress on the root hair shank compared to its tip in the xyloglucan-deficient root hair models is likely the outcome of differential wall thinning in the shank of these models as they elongate **(Fig. 6 B and C)**. This can potentially result in local wall yielding due to lower yield stress of locally thinner walls under the same isotropic turgor and might explain the bulging that is sometimes observed in these root hairs **(Fig. 1A insets).**

### Root hair length remains unaffected by the increasing stiffness of the growth environment in xyloglucan-deficient root hairs, but root hair density decreases

Testing the growth of roots into agar-based media offers a proxy for assessing how crops navigate compacted soils and for evaluating root resilience against mechanical stress. Because agar allows for precise manipulation of mechanical resistance and provides optical clarity, it enables non-destructive, real-time imaging of root architecture (Bello Bello et al., 2025; Xie et al., 2024). These types of studies can also reveal conditional phenotypes that otherwise are not visible by studying root hair growth along the surface of agar plates.

To test whether xyloglucan deficiency influences mechanical responses to media of differing agar concentrations and thus different stiffnesses, we grew Col-0, *xxt1 xxt2*, and *cslc-Δ5* roots in media with agar concentrations of 0.5%, 1%, and 2% **(Fig. 7A)**. We observed that although wild type root hairs grew shorter with increasing media stiffness, as also observed previously (Pereira et al., 2024), the lengths of xyloglucan-deficient root hairs, while shorter overall than wild type, did not vary significantly across agar concentrations **(Fig. 7B)**. Root hair lengths of Col-0 in 0.5%, 1%, and 2% agar media were 240.5 ± 77.52 µm, 135.9 ± 50.79 µm, and 89.80 ± 33.97 µm respectively, whereas *xxt1 xxt2* root hairs were 67.17 ± 20.51 µm, 72.60 ± 28.08 µm, and 67.02 ± 35.90 µm respectively, and *cslc-Δ5* root hairs were 64.88 ± 18.03 µm, 53 ± 24.78 µm, and 58.59 ± 22.37 µm, respectively. Unlike root hair lengths, root hair density decreased with increasing agar concentration in all genotypes **(Fig. 7C)**. Root hair densities of Col-0 in 0.5%, 1%, and 2% agar media were 41.42 ± 11.09, 21.95 ± 17.64, and 5.63 ± 5.05 root hairs per mm of root length, respectively, in *xxt1 xxt2* root hair densities were 23.43 ± 9.61, 8.57 ± 6.40, and 3.05 ± 3.58 root hairs per millimeter of root length, respectively, and in *cslc-Δ5* root hair densities were 17.88 ± 9.84, 4.47 ± 5.56, and 4.37 ± 4.77 root hairs per millimeter of root length, respectively. Root hair densities in both the mutants were significantly lower than wild type in 0.5% and 1% agar media.

**Fig 7.**
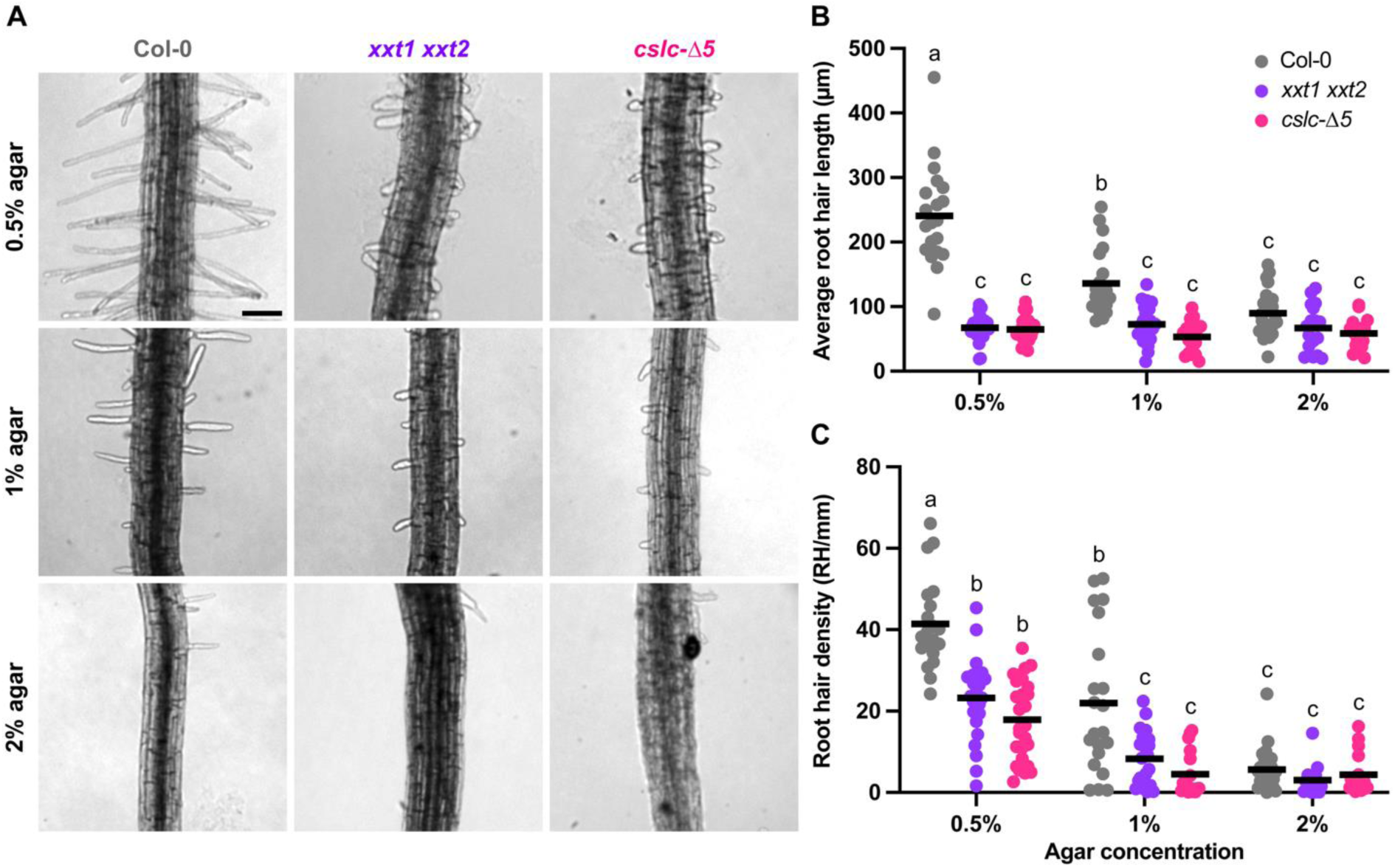
Root hair length but not density remains unaffected in xyloglucan-deficient mutants with increasing media stiffness. **A,** Representative images of Col-0, *xxt1 xxt2*, and *cslc-Δ5* (columns) roots grown in plant media with increase agar concentrations of 0.5%, 1%, and 2% (rows), (Scale bar = 100 µm, applies to all the panels). **B,** Root hair lengths on roots grown in different agar concentrations. Each dot represents the average length of at least five root hairs per seedling. **C,** Graph of root hairs per millimeter of root length. Horizontal lines represent averages. n = 17-27 seedlings per genotype per agar concentration. Letters represent statistical significance, p < 0.05, two-way ANOVA with post-hoc Tukey test.

Decreased root hair density in each genotype **(Fig. 7C)**, was also reflected by decreased root hair count **(Supp. Fig. 6A)**. Similar to the case of root hairs, Col-0 primary roots grew shorter with increasing media stiffness, whereas in both xyloglucan-deficient mutants, root lengths did not change significantly with increasing agar concentrations **(Supp. Fig. 6B)**. In *cslc-Δ5*, roots were shorter than Col-0 at each agar concentration while *xxt1 xxt2* roots did not differ in length from Col-0, unlike what was observed when the roots were grown on plates **(Fig. 1B)**. Root diameters in *xxt1 xxt2* were consistently wider than those in Col-0 irrespective of agar concentrations, whereas *cslc-Δ5* root diameters were only wider than Col-0 in 0.5% agar **(Supp. Fig. 6C)**. Therefore, as stated above, roots and root hairs did show contrasting phenotypes between and within genotypes when grown inside agar media compared to those grown on the surface of media plates. Elucidating the cellular and molecular basis of these phenotypic differences merits detailed exploration but is beyond the scope of this research article.

## Discussion

Arabidopsis root hairs have primary cell walls that are made of cellulose, pectin, and hemicellulosic xyloglucan that that play important roles at different stages of root hair growth (Bernal et al., 2008; Cavalier et al., 2008; Maris et al., 2009; Schoenaers et al., 2024; Zambare & Anderson, 2026). Two proteins from the COBRA-Like family COBL7 and COBL9 localize to the root hair tip in Arabidopsis and control apical cellulose deposition. Root hair growth in the double mutants of *cobl7 cobl9* and double mutants of their rice orthologs *osbc1l1 osbc1l8* terminate prematurely (Li et al., 2022). Another β-1,4-glucan synthase, *At*CSLD3 (KOJAK), a member of cellulose synthase-like (CSL) family, is also required for normal root hair growth as three of its mutant alleles *kjk-2*, *kjk-3*, and *kjk-4,* displayed root hair growth defects since earlier stages (Park et al., 2011; Yang et al., 2020). The chemical status of pectin, whether methyl-esterified or calcium-crosslinked, also governs the biomechanical properties of cell walls (Levesque-Tremblay et al., 2015). Supported by *in vitro* and *in vivo* results, a model suggests that methyl-esterified homogalacturonan is secreted at the root hair tip and is eventually de-esterified by the pectin methylesterases under high pH and gets incorporated into root hair shanks as a ternary complex with LEUCINE-RICH REPEAT EXTENSIN and RAPID ALKALINIZATION FACTOR 22 in the form of concentric rings that potentially restrict radial expansion and promote anisotropic growth (Schoenaers et al., 2024). Periodic oscillations of propidium iodide fluorescence in the root hair tip explain the dynamic change in the chemical status of pectin and suggest that calcium-crosslinked homogalacturonan deposited in root hair shank provides mechanical strength to the shank walls and promotes anisotropic root hair growth (Derbyshire et al., 2007; Rounds et al., 2011). Xyloglucan forms molecular cross-bridges with cellulose and provides flexibility as well as mechanical strength to the cell walls (Chanliaud et al., 2004; Whitney et al., 1999). It is crucial for root hair growth, as xyloglucan-deficient mutants display short and deformed root hairs (Cavalier et al., 2008), but the pathways and mechanisms underlying these phenotypes are unknown. In this study, we explored the cellular and biomechanical properties of two xyloglucan mutants of *Arabidopsis thaliana*, *xxt1 xxt2* and *cslc-Δ5*, that lack xylose sidechains and the glucan backbone of xyloglucan, respectively. Both mutants lack any detectable xyloglucan and display irregular root hair phenotypes (Bou Daher et al., 2024; Cavalier et al., 2008; Kim et al., 2020; Zabotina et al., 2012).

We started by exploring the root hair phenotypes of *xxt1 xxt2* and *cslc-Δ5* in depth. We found that xyloglucan-deficient roots possessed shorter and fewer root hairs that grew slower and were also wider at their shank and bases, than wild type (Col-0) **(Fig 1 and 3)**. These phenotypes suggest more isotropic growth in the root hairs of the mutants, due to potential wall thinning **(Fig. 6 B and C)** in the elongating shank region (Herburger et al., 2022). Higher surface stress in the shank region of root hair FEM can be used as a qualitative proxy for wall thinning **(Fig. 6 B and C, Supp. Table 3).** The decreased anisotropic growth in xyloglucan-deficient root hairs can potentially stem from the altered mechanical properties. To dissect the etiology of these root hair phenotypes, we explored the wall and subcellular properties of root hairs along with their mechanical attributes.

Cell wall thickness and cellulose orientation influence the anisotropic compliance of cell walls and hence anisotropic growth in plant cells (Baskin & Jensen, 2013). We observed that both the *xxt1 xxt2* and *cslc-Δ5* root hairs had thicker cell walls in their shanks with simultaneous wider cross-sections **(Fig. 3)**. This phenotype resembles the characteristics of leaf mesophyll cells of plants grown on NH_4_^+^ versus NO_3_^-^ as a nitrogen source, where cell walls were thinner and thicker respectively and the cell sizes were directly proportional to wall thickness (Podgórska et al., 2017). The walls of etiolated hypocotyls of *xxt1 xxt2* display an increased abundance of homogalacturonan and glucomannan, and hence are thicker (Sowinski et al., 2022), which further corroborates the increased wall thickness observed in root hairs. In another Arabidopsis mutant for xyloglucan galactosyltransferase (*mur3-7*), epidermal cell walls of etiolated hypocotyls, which had shorter cells than wild type, were more porous but thicker compared to those in Col-0, which suggests a compensatory mechanism against compromised wall integrity that might limit cell growth (Xiang et al., 2023). However, defects in cellulose content or organization can also lead to more isotropic growth (Hu et al., 2018; Slabaugh et al., 2014). In our hands, cellulose orientation and optical anisotropy as measured by the FibrilTool plugin did not vary significantly across the three genotypes **(Supp. Fig. S2)**. Therefore, cellulose orientation does not explain the defective growth we observed in xyloglucan-deficient root hairs, although cellulose appeared to be bundled in mutant root hairs **(Supp. Fig. S2)**, similar to what was observed in etiolated hypocotyls of *xxt1 xxt2* (Xiao et al., 2016). This difference in cellulose organization might reflect the absence of xyloglucan as a spacer that prevents cellulose bundling (Eckardt, 2008).

Turgor pressure, besides driving cell expansion, also contributes to the mechanical stiffness of plant cells (Ali et al., 2023). Cell expansion occurs when turgor pressure lies above the yield threshold of the cell wall, but under conditions of compromised wall integrity, turgor pressure decreases, resulting in reductions in cell and organ length (Bou Daher et al., 2024; Qin et al., 2026). Here, we observed that xyloglucan-deficient trichoblasts had lower turgor than wild type **(Fig. 4)**, which provides one explanation for the shorter root hairs observed in these mutants. The epidermal cells of etiolated hypocotyls in *cslc-Δ*5 also showed lower turgor pressure (Bou Daher et al., 2024), which corroborates our results. However, lower turgor pressure in root hairs during growth into stiffer agar was accompanied by faster, but earlier-arresting, root hair elongation in wild-type seedlings (Zambare et al., 2026), implying that a combination of turgor pressure and wall integrity determines growth rate and directionality.

Using the experimentally observed cellular and mechanical parameters in root hairs, we built individual FEMs that recapitulate the characteristic geometries of Col-0, *xxt1 xxt2*, and *cslc-Δ5* root hairs **(Fig. 6A-C)**. Root hair models parameterized with measured values of wall thickness and turgor pressure resulted in modeled growth rates of root hairs that resembled the growth rates observed *in vivo* **(Figs. 1H and 6B)**. Therefore, these models can be used to test the growth dynamics of xyloglucan-deficient root hairs, with some caveats that we highlight later in this section. In our FEMs, we observed non-monotonic stress patterns at the tips of xyloglucan-deficient root hairs. The root hair apex and dome were under lower surface stress compared to their shanks **(Fig. 6)**. This follows the classic hemispherical hoop stress pattern described by the formula σ = P.r/2t, where σ is stress, P is turgor pressure, r is radius, and t is wall thickness. This explains how the geometry of xyloglucan-deficient root hairs distribute surface stress imposed by the turgor pressure which prevents root hair bursting at the tip where the cell wall is being actively assembled and hence is more prone to mechanical failure resulting in root hair rupture (Bernal et al., 2008; Galway et al., 2011). Swapping turgor pressure and cell wall thickness across genotypes revealed that the growth rate and surface stress of root hairs decrease *in silico* with decreasing turgor pressure and with increasing cell wall thickness **(Supp. Tables 1 and 2)**. This further explains the biomechanical mechanisms underlying slower growth in *xxt1 xxt2* and *cslc-Δ*5 root hairs. Our models are limited to the simulation of elastic behavior of the cell wall, as they do not account for the deposition of new wall material and the elasto-viscoplastic behavior of wall during root hair growth (Alline et al., 2026).

In addition to turgor pressure and wall stiffness, root hair growth is mediated by pH, expansin activity, and cytoskeletal dynamics, and several of the latter properties can affect wall mechanics (Bibikova et al., 1998, 1999; Lin et al., 2011; Ringli et al., 2002; Van Bruaene et al., 2004). In *xxt1 xxt2*, petioles are more deformable under tensile stress but show diminished creep under the external application of acidic buffer (pH 4.5) and α-expansins, implying that they are softer but less prone to expansin-mediated creep. We suspect that a lack of expansin-mediated creep in the root hairs of *xxt1 xxt2* and *cslc-Δ5* might result into shorter, slower-growing root hairs. Microtubule stability and orientation are important for root hair initiation and elongation (Sieberer et al., 2005). In *xxt1 xxt2* hypocotyls and meristems microtubule alignment and dynamics are perturbed, and microtubules are less stable and more prone to depolymerization upon application of external mechanical stress (Xiao et al., 2016; Zhao et al., 2019). In xyloglucan-deficient trichoblasts and root hairs, we would expect lower microtubule density, stability, and anisotropy, which might help explain trichoblasts with root hair initiation defects **(Fig. 2C)** and hence fewer **(Fig. 1E)** as well as shorter **(Fig. 1F)** root hairs in *xxt1 xxt2* and *cslc-Δ5*. Actin filaments and actin binding proteins are also important in root hair growth (Ketelaar, 2013). ACTIN2 is essential for bulge site specification and tip growth in root hairs (Ringli et al., 2002). In the etiolated hypocotyls of an Arabidopsis mutant defective in the galactosylation of xyloglucan (*mur3-7*), actin filaments were more sensitive to Latrunculin A treatment, which inhibits actin polymerization (Xiang et al., 2023). Root hair expansion also involves shank-localized cell wall growth as described by pulse-chase labeling experiments that tracked the deposition of galactosylated xyloglucan in the shanks of growing root hairs (Herburger et al., 2022). Xyloglucan galactosyltransferases transfer galactosyl residues on already xylosylated xyloglucan, which is absent in both *xxt1 xxt2* and *cslc-Δ5* root hairs (Zhong et al., 2026). Therefore, actin filaments in trichoblasts and root hairs of xyloglucan-deficient mutants are likely less stable, resulting in fewer and shorter root hairs. These findings highlight the interconnections between cell wall and cytoskeletal stability.

A transcriptomic analysis of differently expressed genes in etiolated seedlings of *xxt1 xxt2* showed downregulation of arabinogalactan protein (*AGP14*) and peroxidase 59 (*PER59*), and upregulation of expansin11 (*EXPA11*) (Ajayi et al., 2024). First two genes function in root hair development, and their downregulation could explain the observed root hair phenotypes **(Fig. 1)** in the xyloglucan deficient mutants. AGP14 controls root hair length, and a loss-of-function mutation of *AGP14* results into abnormally long root hairs both in normal and phosphate-deficient conditions compared to wild type in the corresponding growth conditions (Lin et al., 2011). Therefore, downregulation of *AGP14* in *xxt1 xxt2* might be due to a compensatory response to increase the root hair length. In root hairs, class III peroxidases are critical for crosslinking wall proteins such as extensins via reactive oxygen species (ROS)-mediated pathways and provide mechanical strength to the wall (Marzol et al., 2022). PER59, also sometimes abbreviated as PRX59, expresses in the root hair zone in Arabidopsis (Sundaravelpandian et al., 2013). Downregulation of *PER59* in *xxt1 xxt2* might result in less extensin crosslinking and hence softer cell walls as observed in our Brillouin microscopy data **(Fig. 5) (Table 1)**. PER59 is also important for root hair development in maize and is upregulated when maize seedlings are treated with H_2_O_2_, which is an ROS (Tai et al., 2022). As ROS oscillations modulate root hair growth in Arabidopsis (Monshausen et al., 2007), a downregulation of PER59 in *xxt1 xxt2* might potentially imbalance the critical balance of ROS oscillations and result into shorter root hairs that grow slower **(Fig. 1)**. EXPA11 is a clade-V expansin and is a member of the α-expansin family which also hosts EXPA7 and EXPA18 which are essential for normal root hair growth (Cho & Cosgrove, 2002; Zhou et al., 2026). Complementation of *expa7 expa18* double mutant with EXPA11-mCherry resulted into normal root hair growth (Zhou et al., 2026). Therefore, overexpression of *EXPA11* in *xxt1 xxt2* hints at a compensatory mechanism.

Altogether, the data presented here, along with prior knowledge of how xyloglucan influences cell wall structure and mechanics, point us to a mechanistic framework in which xyloglucan is critical for the developmental decision-making of root epidermal cells to become root hair-forming trichoblasts, for the initiation of root hairs, and for sustaining a rapid rate of root hair elongation during early seedling establishment. This last factor might be a key determinant of seedling anchoring, nutrient acquisition, and survival in disturbed ecosystems such as those favored by Arabidopsis (Choi & Cho, 2019; Lim et al., 2026; Somssich et al., 2016). Thus, the molecular architecture of the cell wall that enables the robust and rapid growth of root hairs can be connected to ecological competitiveness.

## Materials and Methods

### Plants and growth conditions

*Arabidopsis thaliana* (Col-0) and its xyloglucan deficient mutants (*xxt1 xxt2* and *cslc-Δ5*) were used for all the experiments. For incipient plasmolysis and Brillouin microscopy, membrane tagged lines, LTI6b-GFP Col-0, LTI6b-GFP *xxt1 xxt2*, MYF-YFP Col-0, and MYF-YFP *cslc-Δ5* were used. The seeds were first surface sterilized with 30% bleach and 0.1% SDS, washed with sterile Mili-Q^®^ for four times and suspended in 0.15% agar solution for stratification in the dark at 4°C for at least 3-days.

Seedlings were grown on ½ MS media (Caisson labs Murashige and Skoog basal salts MSP01) supplemented with 1% sucrose and 0.8% agar on plates for 5-days before imaging. Lids of the plates were sealed with Micropore^TM^ tape (3M Co.) to facilitate air exchange and avoid contamination. The plate was placed in a plant growth chamber (Percival) at the 24-hour light setting with light intensity 120-150 μmol/m²/s.

#### Root hair growth rate

Col-0, *xxt1 xxt2,* and *cslc-Δ5* seedlings were grown in Ibidi^®^ µ-Slide VI 0.4 Uncoated (Cat.No: 80601). The slides were filled with sterile ½ MS growth media supplemented with 1% sucrose and 0.5% granular agar as described earlier (Zambare et al., 2026). Root hairs growing in the maturation zone were imaged using bright field Z-stack combined with time lapse every minute for a duration of 30 minutes. Movies were analyzed in Fiji (Schindelin et al., 2012).

#### Root hair morphogenetics in different agar concentrations

Seedlings were grown in ½ MS media (Caisson labs Murashige and Skoog basal salts MSP01) supplemented with 1% sucrose and 0.5%, 1%, and 2% agar (Sigma Aldrich A4550, CAS-No:9002-18-0) in 2 ml microcentrifuge tubes (Globe Scientific Inc.), for 5-days before imaging. The seeds were gently sowed beneath the top surface of the media, and 2-3 holes were punched in the cap of microcentrifuge tubes using sterile needles (18G x 1 ½”, AirTite Products Co. Inc.). The punched tubes were covered with Micropore^TM^ tape (3M Co.) to facilitate air exchange and avoid contamination. Tube racks were placed in plant the plant growth chamber (Percival) at the 24-hour light setting with a light intensity of 120-150 μmol/m²/s.

### Confocal microscopy

Zeiss spinning disk confocal microscope (Observer.Z1) with a Yokogawa CSU-X1 spinning head was used for all the experiments except Brillouin microscopy and incipient plasmolysis. Seedlings were mounted on 3x1x1 glass slides (VWR^®^ VistaVision^TM^ Cat.No. 16004-430) in ∼60 µl mounting solution (Mili-Q^®^ or liquid ½ MS media based on the experiment). We made a small chamber using double sided stick-tape (3M^TM^ Scotch^®^ Double Sided Tape 665), and we placed a 24x40 mm cover glass (Corning Cat.No. 2980-244) on the seedling, but it rested on the tape to ensure the seedling was not squished. Zeiss immersion oil (Immersol^TM^ 518 F) was applied to the cover glass when using oil-immersion objectives. We deconvolved confocal images using AutoQuantX2 software.

#### Propidium iodide staining

5-day-old seedlings were incubated in dark for 10 minutes in 10 µg/ml propidium iodide solution (Sigma Aldrich P4864) prepared in Milil-Q^®^. Seedlings were washed twice with Mili-Q^®^ and mounted on glass slide and imaged using 20X objective (0.5 NA) by taking Z-stack series with step distance of 1 µm. 561 nm excitation laser with 617/73 nm emission filter was used to detect propidium iodide fluorescence. These confocal images were used to quantify epidermal cell patterning and length.

#### Pontamine Fast Scarlet 4B (S4B) staining

Five-day-old seedlings were incubated in dark for 30 minutes in 0.01% S4B solution prepared in liquid ½ MS medium as described previously (Anderson et al., 2010). Seedlings were washed once in ½ MS liquid medium and were mounted them on glass slide in ½ MS liquid medium, and imaged under 100X (1.4 NA) objective by taking Z-stack series with step distance of 0.2 µm. S4B stains cellulose and fluoresces when excited with 561 nm laser. A 617/73 nm emission filter was used to detect S4B fluorescence. These confocal images were used to quantify cell wall thickness and cellulose microfibril orientations in root hair shanks.

### Image analysis and quantification

Confocal image analysis was performed with Fiji using both inbuilt tools and plugins (Schindelin et al., 2012). The segmented line tool was used to quantify lengths, and multipoint tool was used to quantify root hairs. Cell wall thickness was measured by plotting the fluorescent intensity profile followed by the Full Width Half Max (FWHM) method (Komis et al., 2015). Epidermal cell patterning was analyzed using BigTrace 0.4.1 plug-in that helps render 3D images for faster cell tracing. The cellulose microfibril orientation was quantified using the FibrilTool plugin (Boudaoud et al., 2014).

### Incipient plasmolysis to calculate turgor pressure

Plasma membrane tagged lines as described earlier in this section were used for these experiments. Five-day-old seedlings were transferred to sorbitol solutions of differing concentrations (0, 0.1, 0.2, 0.3, 0.4, and 0.5 M) for at least 6 mins and subsequently mounted on glass microscope slides (VWR^®^ VistaVision^TM^ Cat.No. 16004-430) with double-sided tape chamber containing sorbitol solution with the respective concentration (∼ 2 mins). Further, the slide was mounted on the microscope stage (Zeiss Axio Imager.M2), and focused for trichoblasts using 40X (1.3 NA) oil immersion objective (∼ 2 mins). Subsequently, the number of plasmolyzed and non-plasmolyzed trichoblasts were counted for the next 5 mins and reported as the percentage plasmolysis and plotted as a function of increasing sorbitol concentration **(Fig. 4A)**. A non-linear regression curve was fitted to find the sorbitol concentration required to plasmolyze 50% of the observed trichoblasts. Corresponding sorbitol concentrations were then entered in the modified universal gas equation ψ = cRT, where ψ is the turgor pressure in Pascals, c is the sorbitol concentration required to plasmolyze 50% of the epidermal cells, R is the universal gas constant (8.3145 J⋅K⁻¹⋅mol⁻¹), and T is the average room temperature of the day in Kelvin. It was important to keep track of the room temperature and follow the time-steps mentioned in brackets as this protocol is time and temperature sensitive.

### Brillouin microscopy

#### Growth conditions

Arabidopsis seeds were sown on 0.5 Murashige Skoog medium with 0.05% MES, 1% sucrose and 1% agar. Plates were stratified for 48 hours at 4 °C and then placed vertically in a Percival growth chamber under long-day conditions (16 hours light/ 8 hours darkness at 22°C, photon density 150 μmol·m^-2^·s^-1^, and 40% humidity).

#### Instrument

The custom-built Brillouin module was described previously (Alonso Baez et al., 2026; Zhang & Scarcelli, 2021). Briefly, an inverted Leica SP8 confocal microscope was coupled to a 2-stage VIPA Brillouin spectrometer. A 532 nm laser (Cobolt) was used as an excitation source and was directed to the backport of the confocal microscope. A 40X (NA 1.1) water immersion objective was used to illuminate the sample, and the light was collected in a backscattering configuration. The scattered light was focused to a single mode optical fiber before entering the spectrometer which consisted of two-stage VIPA etalons with 30 GHz free spectral range (Light Machinery). A qCMOS camera was used to collect the spectra.

#### Imaging and analysis

Image acquisition was set at 100 ms and laser power to 15 mW. Raster imaging with a distance between adjacent pixels of 1 μm (XY plane) and a frame integration of two images was used. Samples were imaged with transmitted light wide-field illumination before and after recording the Brillouin spectra. No phototoxicity effects were observed. A custom MATLAB (Mathworks) script was used to extract the Brillouin peak positions and linewidth after fitting with a Lorenztian function (Alonso Baez et al., 2026). The pixels were ordered according to the stage position to generate heatmaps of frequency shift and linewidth. Water and methanol were regularly measured as reference samples to calibrate the frequency axis. An overlap of the transmitted light and Brillouin images was done manually to define the region of interest (border and area).

#### Moduli calculation

The storage modulus and viscosity (loss modulus) were quantified as described previously (Alonso Baez et al., 2026; Prevedel et al., 2019).

Loss modulus (M’) was calculated as follows,

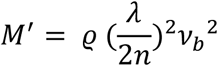

Where, ϱ = mass density of sample, λ = excitation wavelength of the laser (532 nm), n = refractive index of water (1.33), ν_b_ = frequency shift.

The mass density of sample (ϱ) was calculated as follows,

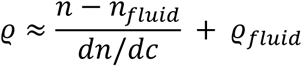

Where, n = refractive index of sample (1.38 on cell wall periphery, 1.35 on root hair area), n_fluid_ = refractive index of water (1.33), dn/dc = refraction increment (dn/dc ∼ 0.145 ml/g for carbohydrates. Used for quantifying mechanical properties at root hair borders. For quantifying mechanical properties in root hair area region, dn/dc ∼ 0.18 ml/g)

Viscosity (ηL) was calculated as follows,

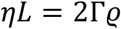

Where, Γ = attenuation coefficient, ϱ = mass density of sample.

Γ was cauclated as follows,

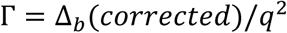

Where, Δ_b_ (corrected) = deconvolved linewidth, q = scattering wavefactor.

Δ_b_ (corrected) was calculated as follows,

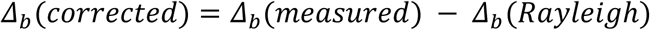

Where, Δ_b_ (Rayleigh) = 0.488 GHz for our setup.

The scattering wavefactor (q) was calculated as follows,

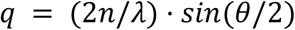

Where, n = refractive index of the sample (1.38 on cell wall periphery, 1.35 on root hair area), λ = excitation wavelength of the laser (532 nm), θ = scattering angle (180° for backscattering).

### Finite Element Modeling

For computational simulations of root hair growth using the Finite Element method, root hairs of each genotype were modeled as hollow cylinders with diameters of 8.72 µm, 15.3 µm, and 15.6 µm for Col-0, *xxt1xxt2*, and *cslc-Δ5* respectively, and capped with half-spheres. Their respective aspect ratios of length:diameter were defined as 15.1, 4.6, and 4.5, respectively. This geometric model was meshed using Gmsh (v 4.15) and imported into Abaqus (v. 2024). The imported root hair mesh was modeled as a shell with an appropriate thickness value reflecting experimental observations of cell wall thickness. The growth of root hair was modeled as a static mechanical response to the uniform turgor pressure applied to the inner surface, which corresponds to the growth of root hairs observed in experiments over 1 min. A linear anisotropic elasticity plane-strain was assigned uniformly across the whole cell wall. The ratio longitudinal:circumferential moduli was assumed to be 1:5 across genotypes because of the similar cellulose orientation observed in their root hair shanks **(Supp. Fig. S4B)**. Genotype specific loading and material drivers such as cell wall thickness and turgor pressure were used to simulate growth of for Col-0, *xxt1 xxt2*, and *cslc-Δ5* root hairs and study surface stress patterns to hypothesize mechanisms of root hair bulging and growth dynamics in xyloglucan-deficient mutants.

### Statistical analyses

We performed statistical analyses with GraphPad v11.0.1. We represented individual data points by dots. Horizontal lines are representative of the mean value. The figure legends below each graph provide information for the sample size, statistical significance annotations (letters), and the statistical tests that we used.

## Acknowledgements

We thank Dr. Firas Bou Daher and Prof. William Gray, University of Minnesota, for providing us with *cslc-Δ5* and MYR-YFP *cslc-Δ5* seeds. Special thanks to the VZ’s thesis committee members at Penn State, Prof. Liana Burghardt, Prof. Gabriele Monshausen, and Prof. Spencer Szczesny for helpful discussions.

## Author contributions

VZ and CTA conceived the ideas and designed experiments, VZ, GM, LAB performed experiments and analyzed data, VZ and CTA interpreted data, HY and VZ built and tested the FEM, VZ and GM drafted the manuscript, CTA and TH acquired funding and supervised the project, and all authors reviewed, edited, and approved the manuscript.

## Data availability

The data underlying this article will be shared on reasonable request to the corresponding author.

