## Supplementary material for "Mechanisms underlying the defective development of xyloglucan-deficient root hairs in *Arabidopsis thaliana*"

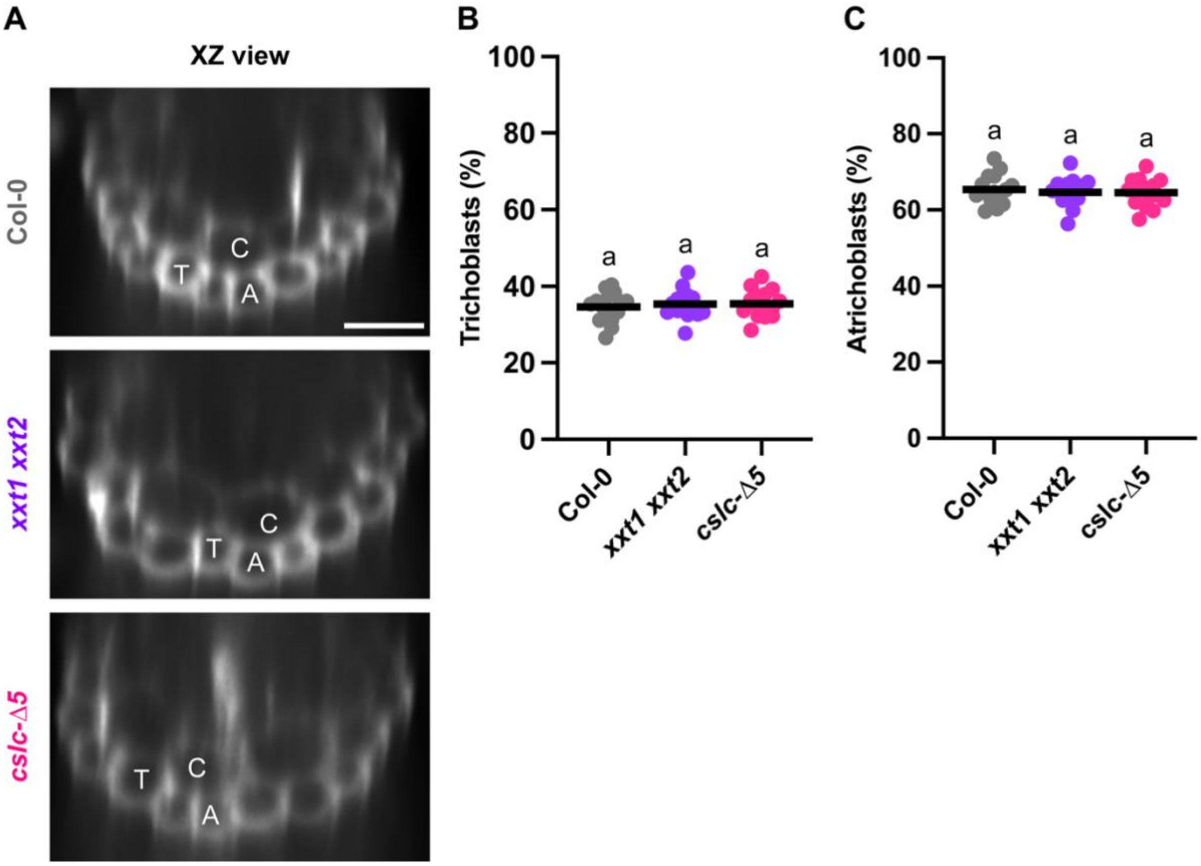


**Supplementary Fig 1. Xyloglucan deficiency does not affect epidermal cell patterning in roots**

**A,** Images represent transverse optical cross-sections of 5-day old roots of Col-0, *xxt1 xxt2*, and *cslc*-Δ*5* grown on media plates (Scale Bar = 50 μm, applies to all the panels). Fluorescence signal represents propidium iodide staining. C = Cortical cell, T = Trichoblast, A= Atrichoblast. **B,** Percentage of root epidermal cells that are trichoblasts. **C,** Percentage of root epidermal cells that are atrichoblasts. Cells were observed and numbers were averaged throughout the root maturation zone. Only the cells from the observable half of the root were scored. Each point on the graph is one seedling (n = 15 seedlings per genotype from three independent experiments). Horizontal lines represent averages. Different letters above the graph represent if a significant difference between the groups is present, $p < 0.05$, one-way ANOVA and post-hoc Tukey test.

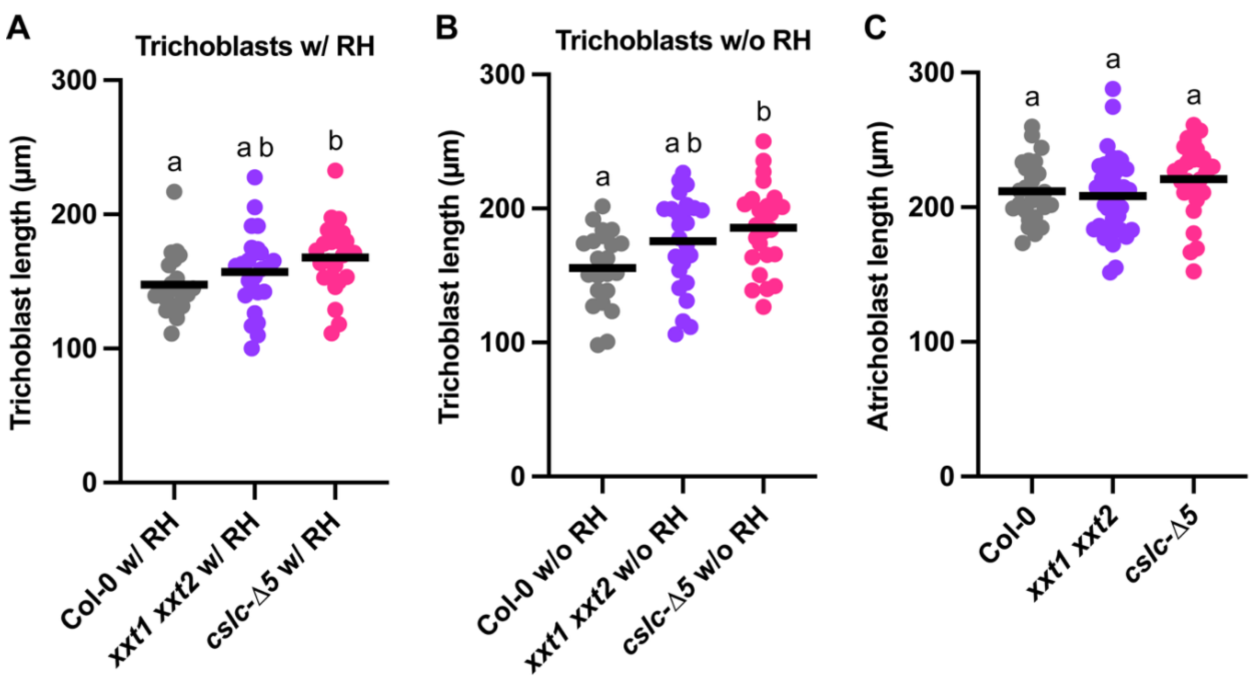


**Supplementary Fig 2. Xyloflucan-deficient trichoblasts are longer than wild type whereas atrichoblast lengths do not vary significantly across genotypes**

**A,** Lengths of trichoblasts from the root maturation zone that have a root hair. **B,** Lengths of trichoblasts from the root maturation zone that do not have a root hair, (n = 22-25 seedlings per genotype from five independent experiments). **C,** Lengths of atrichoblasts from the root maturation zone (n = 28-36 seedlings per genotype from six independent experiments). Horizontal lines represent averages. Different letters above the graph represent if a significant difference between the groups is present, $p < 0.05$, one-way ANOVA and post-hoc Tukey test.

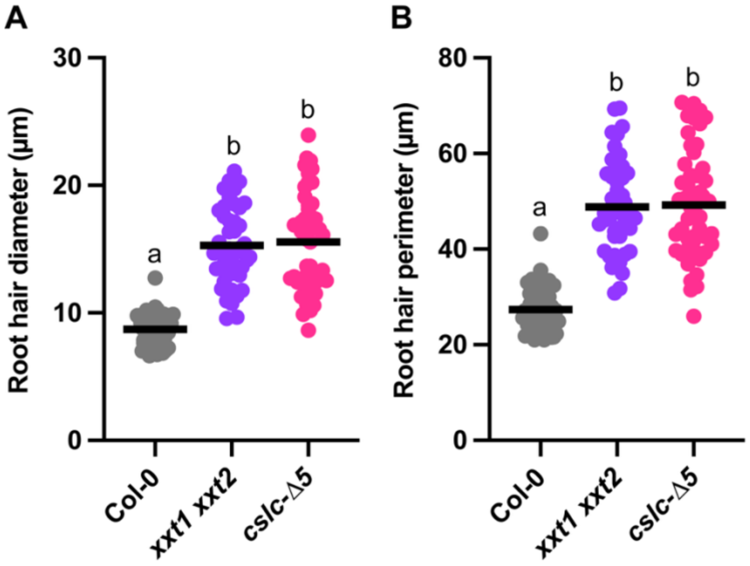


**Supplementary Fig 3. Xyloflucan-deficient root hairs are wider than wild type**

**A,** Average root hair diameter along the root hair shank. **B,** Average root hair perimeter along the root hair shank. (n = 45-54 root hairs per genotype from at least 15 seedlings). Horizontal lines represent averages. Different letters above the graph represent if a significant difference between the groups is present, $p < 0.05$, one-way ANOVA and post-hoc Tukey test.

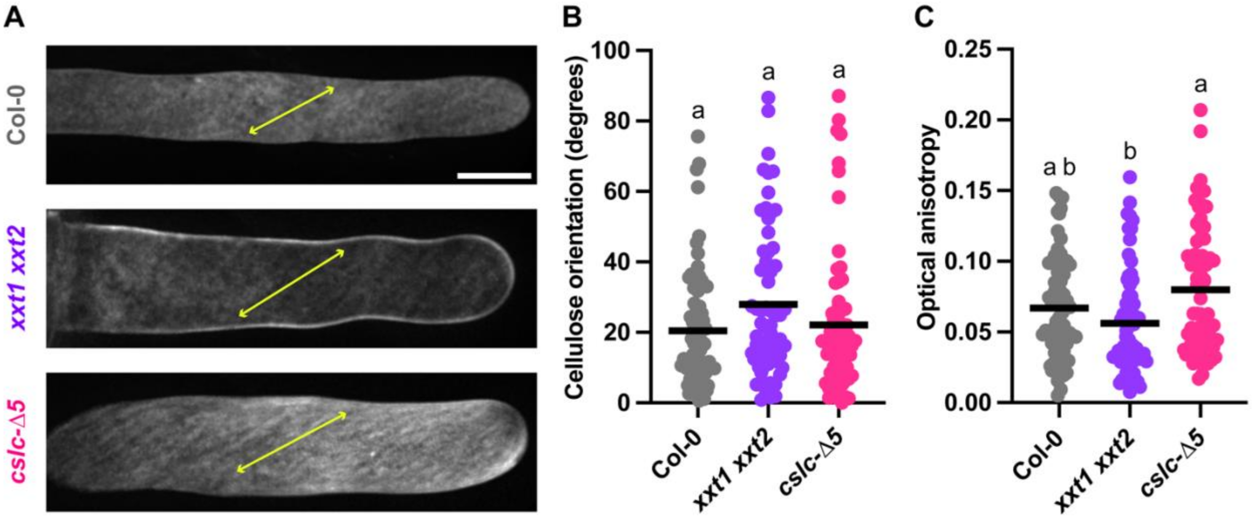


**Supplementary Fig 4. Xyloglucan deficiency has no effect on cellulose orientation in root hairs.**

**A,** Images represent 5-day old root hairs of Col-0, *xxt1 xxt2*, and *cslc*-Δ*5* grown on media plates (Scale Bar = 10 µm, applies to all the panels). Fluorescence signal represents S4B stain that specifically bind to cellulose fibrils. Double headed arrows represent average cellulose orientation. **B,** Cellulose orientation in root hair shanks. **E,** Optical anisotropy as quantified by FibrilTool. Each point represents one root hair (n = 64 - 70 root hairs per genotype). Horizontal lines represent averages. Different letters above the graph represent statistically significant difference, $p < 0.05$, one-way ANOVA and post-hoc Tukey test.

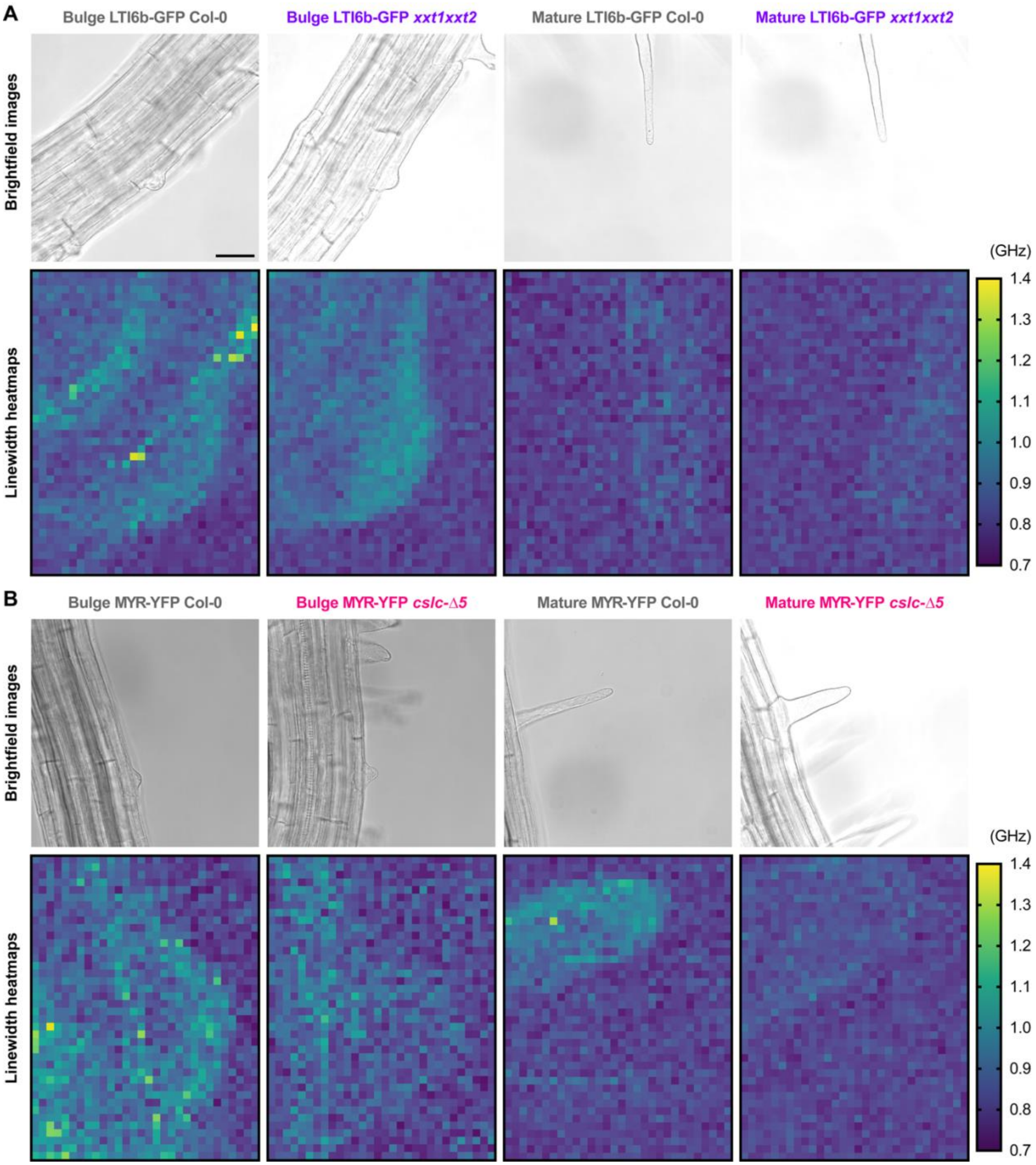


**Supplementary Fig 5. Brillouin microscopy reveals no change in the viscosity of root hairs that are xyloglucan-deficient compared to wild type.**

**A, Upper row:** Brightfield images of root hair bulges and mature root hairs of Col-0 and *xxt1 xxt2* (Scale bar = 100 μm, applies to all the confocal panels), **Lower row:** Representative heatmaps showing linewidth values of the Brillouin spectra acquired from corresponding root hairs. **B, Upper row:** Brightfield images of root hair bulges and mature root hairs of Col-0 and *cslc*-Δ*5*, **Lower row:** Representative heatmaps showing linewidth values of the Brillouin spectra acquired from corresponding root hairs.

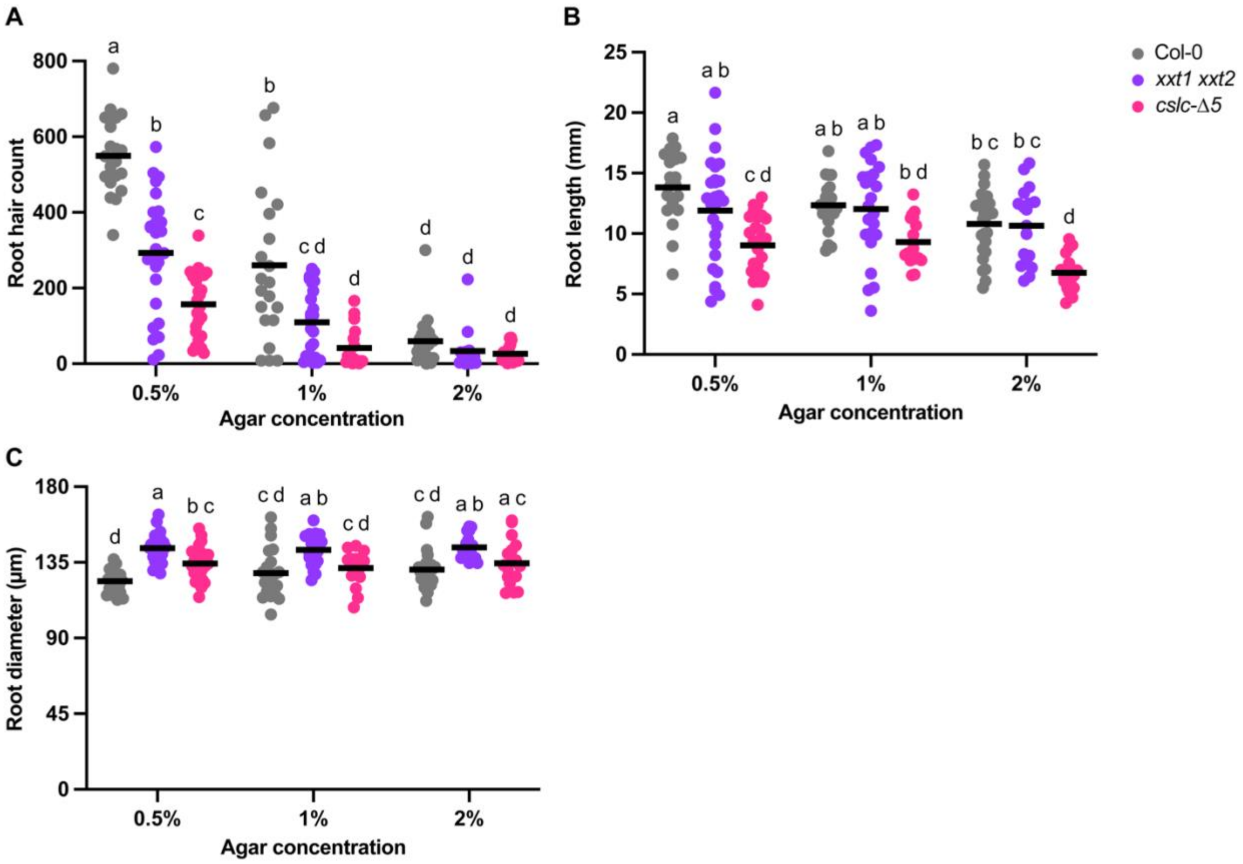


**Supplementary Fig 6. Morphological analysis of root hairs and primary roots under differing media stiffness in wild-type and xyloglucan-deficient seedlings.**

**A,** Number of root hairs per seedling in each genotype grown at different agar concentrations. **B,** Lengths of roots grown in media with different agar concentrations. **C,** Diameters of roots grown in media with different agar concentrations. n = 17-27 seedlings per genotype per agar concentration. Horizontal lines represent averages. Letters represent statistical significance, $p < 0.05$, two-way ANOVA with post-hoc Tukey test.

**Supplementary Table 1: Xyloglucan-deficient root hairs show overall lower surface stress than wild type**

The values below show a detailed description for Fig. 6 E-G.

| Tip stress (MPa) | | | Dome stress (MPa) | | | Subapical stress (MPa) | | |
|---|---|---|---|---|---|---|---|---|
| Col-0 | *xxt1 xxt2* | *cslc-Δ5* | Col-0 | *xxt1 xxt2* | *cslc-Δ5* | Col-0 | *xxt1 xxt2* | *cslc-Δ5* |
| 52.01 | 5.10 | 3.27 | 56.46 | 9.16 | 7.52 | 54.23 | 10.19 | 8.58 |
| 43.72 | 3.95 | 1.17 | 47.43 | 7.76 | 4.12 | 45.57 | 9.70 | 4.86 |
| 22.46 | 4.36 | 3.01 | 24.45 | 8.24 | 7.15 | 24.45 | 9.21 | 8.18 |
| 376.05 | 5.61 | 4.86 | 376.05 | 9.81 | 9.72 | 376.05 | 10.86 | 10.93 |
| 34.27 | 5.97 | 3.62 | 35.71 | 10.25 | 8.02 | 34.27 | 11.32 | 9.12 |
| 90.51 | 5.70 | 4.53 | 90.51 | 9.92 | 9.28 | 90.51 | 10.98 | 10.46 |

**Supplementary Table 2: Finite element results after swapping turgor pressure**

Average turgor pressure values of different genotypes Col-0 (0.798 MPa), *xxt1 xxt2* (0.637 MPa), and *cslc*-Δ*5* (0.605 MPa), were swapped with every other genotype to run finite element models. Average turgor pressure value for Col-0 was obtained by averaging the mean turgor pressures of LTI6b-GFP Col-0 and MYR-YFP Col-0 (Fig. 4B).

| | **Input** | | **Output** | | | |
|---|---|---|---|---|---|---|
| Genotype | Cell wall thickness (nm) | **Turgor pressure (MPa)** | Growth rate (μm/min) | Tip stress (MPa) | Dome stress (MPa) | Subapical stress (MPa) |
| Col-0 | 466 | 0.798 | 1.26 | 52.01 | 56.46 | 54.23 |
| Col-0 | 466 | **0.637** | 0.68 | 4.59 | 7.80 | 8.60 |
| Col-0 | 466 | **0.605** | 0.50 | 4.36 | 6.68 | 7.26 |
| *xxt1 xxt2* | 548 | **0.798** | 0.71 | 7.62 | 13.02 | 13.85 |
| *xxt1 xxt2* | 548 | 0.637 | 0.57 | 5.10 | 9.16 | 10.19 |
| *xxt1 xxt2* | 548 | **0.605** | 0.54 | 4.58 | 8.52 | 9.51 |
| *cslc*-Δ*5* | 572 | **0.798** | 0.77 | 5.97 | 11.17 | 12.47 |
| *cslc*-Δ*5* | 572 | **0.637** | 0.62 | 3.66 | 8.07 | 9.18 |
| *cslc*-Δ*5* | 572 | 0.605 | 0.59 | 3.27 | 7.52 | 8.58 |

**Supplementary Table 3: Finite element results after swapping cell wall thickness**

Average cell wall thicknesses of different genotypes Col-0 (466 nm), *xxt1 xxt2* (548 nm), and *cslc-Δ5* (572 nm), were swapped with every other genotype to run finite element models.

| | Input | | Output | | | |
|---|---|---|---|---|---|---|
| Genotype | Turgor pressure (MPa) | **Cell wall thickness (nm)** | Growth rate (µm/min) | Tip stress (MPa) | Dome stress (MPa) | Subapical stress (MPa) |
| Col-0 | 0.798 | 466 | 1.26 | 52.01 | 56.46 | 54.23 |
| Col-0 | 0.798 | **548** | 0.93 | 6.91 | 10.97 | 11.99 |
| Col-0 | 0.798 | **572** | 0.74 | 4.99 | 8.43 | 9.29 |
| *xxt1 xxt2* | 0.637 | **466** | 0.67 | 7.18 | 11.67 | 12.79 |
| *xxt1 xxt2* | 0.637 | 548 | 0.57 | 5.10 | 9.16 | 10.19 |
| *xxt1 xxt2* | 0.637 | **572** | 0.54 | 4.66 | 8.63 | 12.25 |
| *cslc-Δ5* | 0.605 | **466** | 0.72 | 5.17 | 10.05 | 11.27 |
| *cslc-Δ5* | 0.605 | **548** | 0.62 | 3.59 | 7.97 | 9.07 |
| *cslc-Δ5* | 0.605 | 572 | 0.59 | 3.27 | 7.52 | 8.58 |
